# EEG source imaging technique to investigate sleep oscillations for simultaneous EEG-fMRI

**DOI:** 10.1101/2024.08.13.607843

**Authors:** Makoto Uji, Aude Jegou, Nathan E. Cross, Florence B. Pomares, Aurore A. Perrault, Alex Nguyen, Umit Aydin, Kangjoo Lee, Chifaou Abdallah, Birgit Frauscher, Jean-Marc Lina, Thien Thanh Dang-Vu, Christophe Grova

## Abstract

Simultaneous recording of electroencephalography (EEG) and functional magnetic resonance imaging (fMRI) is a widely used non-invasive neuroimaging technique in sleep studies. However, EEG data are strongly influenced by two types of MRI-related artefacts: gradient artefacts (GA) and ballistocardiogram artefacts (BCG). If artefacts correction is suboptimal, the BCG obscures the EEG signals below 20Hz and could make it difficult to investigate sleep oscillations, especially sleep spindles, sleep specific oscillations occurring within 11-16Hz frequency band. We previously demonstrated the utility of beamforming spatial filtering in correcting MRI-related artefacts on EEG. Here, we investigated the use of beamforming spatial filtering for improving the detection of EEG oscillations during sleep, in particular, 1) to accurately estimate single-event spindle EEG power changes, and 2) to demonstrate the potential improvement of fMRI general linear model (GLM) analysis when involving such additional EEG information. We analysed EEG-fMRI data acquired during a recovery nap after sleep deprivation in 20 young healthy participants (12 females, 8 males, age = 21.3 ± 2.5 years). Based on spindle events (onset and duration) detected by trained sleep scorers on BCG corrected EEG signals through a conventional average artefact subtraction (AAS) method, we compared four different EEG processing steps: non-BCG corrected; AAS BCG corrected; beamforming BCG corrected; beamforming+AAS BCG corrected. These processing steps consist of non-BCG corrected and AAS BCG corrected considered either at the sensor level or at the source-level (after beamformer localization) to evaluate the impact of the BCG artefact on the detection of spindle activity. Then we further investigated four different fMRI GLM approaches using 1) the spindle onset and duration (GLM1), 2) spindle onset, duration, and parametric modulation of single-spindle power change from the Cz electrode of the AAS BCG corrected data (GLM2), 3) spindle onset, duration, and parametric modulation of single-spindle power change from the beamforming+AAS BCG corrected (GLM3) and 4) spindle onset, duration, and parametric modulation of single-spindle power change from the beamforming BCG corrected data (GLM4). We found that the beamforming approach did not only attenuate the BCG artefacts, but also recovered sleep spindle activity occurring during NREM sleep. Furthermore, this beamforming approach allowed us to accurately estimate single-event power change of each spindle in the source space when compared to the channel level analysis, and therefore to further improve the specificity of fMRI GLM analysis, better localizing the recruited brain regions during spindles. Our findings show the benefit of applying beamforming source imaging technique to EEG-fMRI acquired during sleep. We demonstrate that this approach would be beneficial especially for long EEG-fMRI data acquisitions (i.e., sleep, resting-state), when the BCG correction becomes problematic due to inherent dynamic changes of heart rates. Our findings extend previous work regarding the application of the source imaging to the sleep EEG-fMRI. Combining with this advanced methodology and analysis, sleep EEG-fMRI will help us better understand the functional roles of human sleep.

## Introduction

Sleep is crucial for both cognition, such as memory and learning (Diekelmann and Born, 2010; Fogel et al., 2012), and maintenance of healthy brain function (Eide et al., 2021; Fultz et al., 2019). Simultaneous recording of electroencephalography (EEG) and functional magnetic resonance imaging (fMRI) can provide greater specificity and sensitivity of neural activities by combining the complementary advantages of each method (high temporal resolution of EEG & high spatial resolution of fMRI) when compared to a single neuroimaging method (EEG or fMRI alone) (Dang-Vu et al., 2010b; Duyn, 2012; Picchioni et al., 2013). Thus, this multimodal method is a widely used non-invasive neuroimaging technique that can be considered for sleep studies. EEG-fMRI has been used to investigate the functional roles of sleep related EEG activity, such as sleep spindles (Bergmann et al., 2012; Caporro et al., 2012; Dang-Vu et al., 2011; Fogel et al., 2017; Jegou et al., 2019; Schabus et al., 2007). Sleep spindles are transient bursts of neural activity with a typical frequency range of 11-16Hz, that last between 0.5 and 3 seconds. They predominantly occur during non-rapid-eye movement (NREM) sleep stage 2 but can also occur during NREM3 (Antony et al., 2019; De Gennaro and Ferrara, 2003; Fernandez and Lüthi, 2020). Since spindles appear to have several different functional roles, previous EEG-fMRI findings demonstrated that sleep spindles are expected to play a key role in sleep-dependent memory processing (Bergmann et al., 2012; Fogel et al., 2017; Jegou et al., 2019) and also selective processing and suppressing of external stimuli, such as auditory tones, during sleep to maintain sleep and ensure a good quality of sleep (Dang-Vu et al., 2011; Schabus et al., 2012).

Although EEG-fMRI has been widely used to advance our understanding of the human sleeping brain, EEG data are strongly influenced by two types of MRI related artefacts, known as gradient artefacts (GA) and ballistocardiogram artefacts (BCG) (Allen et al., 2000, 1998; Niazy et al., 2005). The GA artefact is induced by temporally varying magnetic field gradients used for MR imaging, whereas the BCG artefact is produced by cardiac pulse driven tiny head motions in the strong magnetic field of the MRI scanner (Debener et al., 2008; Mullinger and Bowtell, 2011). Compared to GA, the BCG artefacts are more challenging to cope with due to their inherent variability and dynamic changes over time (Laufs et al., 2008). The BCG particularly obscures the EEG signals below 20Hz, and could make it difficult to investigate sleep oscillations such as sleep spindles (11-16Hz) if the correction is suboptimal (Bonmassar et al., 2002; Uji et al., 2021; Xia et al., 2014, 2013). Therefore, for EEG-fMRI data acquired during sleep, the BCG artefacts need to be carefully removed, and a reliable BCG correction method needs to be developed and established.

Among several different approaches that have been proposed to correct BCG artefacts, the most used method is average artefact subtraction (AAS), which requires detecting the cardiac pulses (R-peak) from simultaneous electrocardiograph (ECG) recording with high precision to construct averaged EEG BCG artefact templates that will be later subtracted from the EEG data (Abreu et al., 2018; Allen et al., 2000, 1998; Bullock et al., 2021). However, the ECG signal in the MRI scanner is often distorted, making automatic detection of R-peaks problematic (Chia et al., 2000; Mullinger et al., 2008b), although a few research groups have attempted to use facial and temporal electrodes of high-density EEG for the BCG detections (Iannotti et al., 2015). Thus, accurate BCG correction often requires manual correction of R-peaks detections which is time-consuming especially when considering a long recording required for sleep studies (i.e., maximum up to 2 hours). In addition, heart rates during sleep recording vary across different vigilance stages (Tobaldini et al., 2013), therefore impacting the accuracy of BCG correction approaches. Thus, detecting R-peak events from the ECG is difficult and often become unreliable.

In this study, we propose an alternative BCG correction method to the AAS approach: beamforming spatial filtering approach to attenuate the BCG artefacts (Brookes et al., 2008a; Uji et al., 2021), assessing for the first time its relevance when applied on sleep EEG-fMRI data. This approach appears as a promising denoising technique, and has been only recently considered in the context of EEG-fMRI studies (Brookes et al., 2009, 2008a; Mullinger and Bowtell, 2011; Uji et al., 2021). Especially, the beamforming technique is a well known highly efficient method to attenuate artefactual signals which have different spatial origins from the underlying signal of interest, such as eye movements (Cheyne et al., 2006) or orthodontic metal braces (Cheyne et al., 2007) in MEG, and the MRI-related artefacts of EEG signal in EEG-fMRI (Brookes et al., 2008a; Uji et al., 2021).

The development of an advanced beamforming technique in EEG-fMRI can further help to improve the sensitivity and specificity of fMRI blood oxygen level-dependent (BOLD) analysis that use information from concurrent EEG recordings (Ikemoto et al., 2022; Jorge et al., 2014; Laufs, 2012; Moeller et al., 2020; Murta et al., 2015; Philiastides et al., 2021). In such approaches, EEG can be used as a temporal marker of neuronal activity or to quantify specific properties of the EEG response (i.e., spindles), and fMRI is then used to study the corresponding hemodynamic response elicited by such neuronal activity within the whole brain. The most common method of the EEG-informed fMRI is General Linear Method (GLM) (Friston et al., 1998), where events or signals of interest detected on EEG are used to examine hemodynamic BOLD signal changes acquired with fMRI. In general, a consistent fixed binary boxcar regressor (i.e., 0 or 1) based on EEG time traces is convolved with a hemodynamic response function (HRF) for the fMRI GLM analysis. However, previous findings demonstrated that the amplitude and variability of BOLD responses could be better explained by single-trial or event variability from EEG signals as compared to the conventional GLM of consistent amplitude responses (LeVan et al., 2010; Mayhew and Bagshaw, 2017; Mullinger et al., 2014; Vulliemoz et al., 2010). This single event variability is also present in sleep spindles in terms of its power or amplitude even from the same electrode at different NREM stages (N2 vs. N3) (Cox et al., 2017). Furthermore, the spindle variability in terms of the peak frequency, amplitude, and duration within the same or across brain regions has recently caught more attention to better understand its underlying mechanism (Frauscher et al., 2015; Mölle et al., 2011; von Ellenrieder et al., 2020; Zerouali et al., 2014). Thus, modelling the spindle related BOLD signals with a conventional boxcar ([1 0]) hemodynamic convolution in fMRI GLM analysis might be too simplistic. Although this conventional approach has been successful to examine spindles (Dang-Vu et al., 2011; Fogel et al., 2017; Jegou et al., 2019; Schabus et al., 2012, 2007), a more accurate approach would be to add such single-spindle variability into the GLM model to improve sensitivity and specificity of the fMRI GLM analysis in sleep studies, as this was demonstrated in epilepsy (Grouiller et al., 2011; LeVan et al., 2010; Vulliemoz et al., 2010) and cognitive and sensory tasks (Mullinger et al., 2017, 2014, 2013; Pakenham et al., 2020; Uji et al., 2018; Wilson et al., 2019). To this end, we propose to use EEG source imaging technique (beamforming approach) to estimate the single-event source signal power variability more accurately for the parameterized fMRI GLM analysis to improve its sensitivity and specificity.

The aim of this study was 1) to investigate the use of a source imaging technique (beamforming spatial filtering) to attenuate the BCG artefacts in EEG-fMRI acquired during sleep and to accurately estimate single-event sleep spindle EEG power changes, and 2) to examine if the proposed techniques can improve the fMRI GLM analysis implementing such EEG information. To achieve this goal, we analysed the EEG-fMRI data acquired during a recovery nap after sleep deprivation in 20 young healthy participants. Based on spindle events (onset and duration) detected by trained sleep scorers when reviewing the BCG corrected EEG signals through a conventional AAS method, we first compared four different EEG processing steps (non-BCG correction; AAS-based BCG correction; beamforming BCG correction; beamforming+AAS-based BCG correction). These processing steps consist of considering non-BCG corrected and AAS BCG corrected either at the sensor level or at the source-level to examine the impact of BCG correction strategies on the assessment of spindle activity. Then we further investigated four different fMRI GLM approaches using 1) the spindle onset and duration (GLM1), 2) spindle onset, duration, and parametric modulation of single-spindle power change from the Cz electrode, after applying AAS BCG correction (GLM2), 3) spindle onset, duration, and parametric modulation of single-spindle power change from the beamforming+AAS BCG corrected data (GLM3), and 4) spindle onset, duration, and parametric modulation of single-spindle power change from the beamforming BCG corrected data (GLM4).

## Methods

The EEG-fMRI data were acquired using a protocol previously published in Cross et al. (2021a), and reused for our specific aim to investigate the use of the beamforming spatial filtering technique in simultaneous EEG-fMRI acquired during sleep. The original study aimed at examining the effect of sleep deprivation and a subsequent recovery nap on functional connectivity patterns across cognitive tasks of attention, vigilance, and working memory. Details of the study designs and findings can be found in (Cross et al., 2021a, 2021b).

20 young healthy participants (12 females, 8 males, age = 21.3 ± 2.5 years) took part in the study and were screened for the absence of sleep disorders (insomnia, sleep apnea syndrome, hypersomnolence, restless legs syndrome, parasomnias), neurological or psychiatric conditions (e.g., epilepsy, migraine, stroke, chronic pain, major depression, anxiety disorder, psychotic disorder) and current use of psychotropic medications or cannabis. All subjects provided written informed consent prior to the start of the study that was approved by the Central Research Ethics Committee of the Quebec Ministry of Health and Social Services. More details about the participants’ characteristics can be found in Cross et al. (2021a). Since the aim of this study was to examine the use of the beamforming spatial filtering technique in simultaneous EEG-fMRI acquired during sleep, 60-minute recovery nap data following sleep deprivation from all 20 healthy participants were pre-processed and analysed.

### Simultaneous EEG-fMRI data acquisition during sleep

#### MRI data acquisition

MRI data were acquired with a 3T GE scanner (General Electric Medical Systems, Wisconsin, US) using an 8-channel head coil at the PERFORM Centre of Concordia University. Functional MRI data were acquired using a gradient-echo echo-planar imaging sequence (TR = 2500ms, TE = 26ms, FA = 90°, 41 transverse slices, 4-mm slice thickness with a 0% inter-slice gap, FOV = 192 x 192 mm, voxel size = 4×4×4mm, matrix size = 64 × 64, maximum 1440 volumes). High-resolution T1-weighted anatomical MR images were acquired using a 3D BRAVO sequence (TR = 7908 ms, TI = 450 ms, TE = 3.06 ms, FA = 12°, 200 slices, voxel size = 1.0×1.0×1.0 mm, FOV = 256 × 256 mm). During all EEG-fMRI sessions, the Helium compression pumps used for cooling down MR components were switched off to reduce MR environment related artefacts infiltrating the EEG signal (Mullinger et al., 2008a; Rothlübbers et al., 2015). To minimize movement-related artefacts during the scan, MRI-compatible foam cushions were used to fix the participant’s head in the MRI head coil (Mullinger and Bowtell, 2011).

#### EEG data acquisition

EEG data were acquired using an MR compatible 256 high-density geodesic sensor EEG array (Electrical Geodesics Inc (EGI), Magstim-EGI, Oregon USA). The EEG cap included 256 sponge electrodes referenced to Cz that covered the entire scalp and part of the face. EEG data were recorded using a battery-powered MR-compatible amplifier shielded from the MR environment that was placed next to the participant inside the scanning room. The impedance of the electrodes was initially maintained below 20kΩ and kept to a maximum of 70kΩ throughout the recording. Data were sampled at 1000Hz and transferred outside the scanner room through fibre-optic cables to a computer running the Netstation software (v5, EGI). The recording of EEG was phase-synchronized to the MR scanner clock (Sync Clock box, EGI), and the onset of every scanner repetition time (TR) period was recorded in the EEG data to facilitate MR gradient artefact correction. ECG was also collected via two MR compatible electrodes placed between the 5^th^ and 7^th^ ribs and above the heart close to the sternum and recorded at 1000Hz through a bipolar amplifier (Physiobox, EGI). Following EEG-fMRI data acquisition, EEG electrode locations were digitized using the EGI Geodesic Photogrammetry System (GPS) to facilitate individualized co-registration of electrode positions with each participant’s anatomical image (Russell et al., 2005).

### Data analysis

#### EEG data analysis

For both EEG and ECG data, gradient artefacts (GA) were first corrected in BrainVision Analyzer2 (Version 2.2.0, Brain Products GmbH, Gilching, Germany) with the standard average artefact subtraction (AAS) method. The AAS method used sliding window templates formed from the averages of 21 TRs, which were subtracted from each occurrence of the respective artefacts for each electrode (Allen et al., 2000; Wilson et al., 2019). GA corrected data were subsequently bandpass filtered (EEG: 1-100 Hz with notch filter of 60Hz), and downsampled (500 Hz). Following the GA correction, cardiac R-peaks were automatically detected from the ECG recording in BrainVision Analyzer2, checked visually and corrected manually if necessary. These R-peak events were used to inform ballistocardiogram (BCG) correction.

From the same GA corrected datasets, two different datasets for each subject were created: non-BCG corrected, and BCG corrected data. The AAS BCG artefact correction was performed using the standard sliding template average-artefact subtraction considering 11 R-peak events per template, as implemented in BrainVision Analyzer2 (Mullinger et al., 2014).

For both non-BCG corrected and BCG corrected data, two lowest rows of electrodes around the neck and face (69 electrodes) were removed from EEG leaving 187 electrodes for further analysis (Supplementary Figure S2) (Hedrich et al., 2017). From the remaining 187 electrodes, noisy and bad electrodes were identified as the ones exceeding 5 standard deviations above the normalized time-course (Delorme and Makeig, 2004; Uji et al., 2019) and were then interpolated using neighbour electrodes. Resulting data were then further downsampled to 250Hz and re-referenced to an average of all non-noisy channels using EEGLAB software (https://sccn.ucsd.edu/eeglab) (Delorme and Makeig, 2004). After re-referencing, all timepoints before and after the MRI scanning were removed, and further analyses were conducted on EEG data acquired during MRI scanning.

#### Sleep staging

Sleep staging was conducted on the AAS BCG corrected data following the standard criteria (Iber et al., 2007). Prior to sleep staging, the BCG corrected data was further band-pass filtered between 0.5 and 20 Hz to remove low-frequency drift and high-frequency noise, and re-referenced to the linked mastoids. Scoring of the sleep stages was performed in conjunction by two trained scorers (NEC & AAP) using the Wonambi toolbox (https://github.com/wonambi-python/wonambi) (using F3, Fz, F4, C3, C4, O1, O2 channels), in order to obtain measures of sleep including total sleep time and the duration of sleep stages (Table 1).

**Table 1.** Mean (±SD) sleep architecture and spindle characteristics during the recovery nap.

|  | **Mean** | **SD** |
| --- | --- | --- |
| Sleep latency (min) | 5.32 | 9.77 |
| Total sleep duration (min) | 46.60 | 14.48 |
| Sleep efficiency (%) | 87.22 | 12.38 |
| NREM1 (%<br>N=20) | 6.59 | 5.20 |
| NREM2 (%<br>N=20) | 70.41 | 19.51 |
| NREM3 (%<br>N=16) | 23.00 | 21.16 |
| Sleep fragmentation (n/h) | 8.91 | 5.62 |
| Latency to NREM3 (min) | 15.91 | 5.16 |
| Number of spindles (n) | 168.65 | 70.10 |
| Spindle density (n/min) | 4.23 | 2.34 |
| Spindle duration (s) | 0.83 | 0.05 |

#### Spindle detections

Spindles were detected using a two-step semi-automatic procedure. First, FASST algorithm (fMRI Artifact rejection and Sleep Scoring Toolbox) developed by Cyclotron Research Centre, University of Liège, Belgium, was used to automatically detect the spindles (Dang-Vu et al., 2011; Jegou et al., 2019; Leclercq et al., 2011; Schabus et al., 2007). The FASST algorithm was applied on the F3, Fz, F4, C3, C4, P3, Pz, and P4 during N2 and N3 stages following Mölle’s approach (Mölle et al., 2002). Specifically, these EEG signals were filtered in spindle frequency band (11-16Hz). Then, the root mean square (RMS) of the filtered signals was calculated during the N2 stages, and each electrode threshold was determined corresponding to 95th percentile of the N2 stage amplitudes. The RMS in the alpha band (8-12Hz) (RMSa) and in the muscle artefact band (30-40Hz) (RMSm) were also calculated in order to minimize false detections (Schimicek et al., 1994). Spindles were detected automatically if they were above the 95% threshold and the ratio between RMSa and the RMS in the spindle band (RMSs) was equal or below 1.2 (RMSa/RMSs≤1.2) and RMSm < 5µV.

Second, four trained scorers (NEC, AAP, CA, BF) reviewed and corrected the automatically detected spindles following the standard guideline (Iber et al., 2007). Data from each participant were reviewed by at least two scorers who accepted or rejected the automatically detected spindles from the FASST algorithm. Additionally, on selected clean 5min epochs in both N2 and N3 stages (i.e., validated NREM period), the scorers were allowed to add spindles if the FASST algorithm failed to detect it. The final outcomes of the spindle detections were determined by the consensus between the two reviewers.

#### EEG beamforming

A scalar linearly constrained minimum variance (LCMV) beamforming analysis (Brookes et al., 2008a; Uji et al., 2018; 2021; van Veen et al., 1997) with a regularization parameter of 0.01% was employed. Individual boundary element method (BEM) head models (Brookes et al., 2008b; Litvak et al., 2010) was used to reconstruct broadband (1-100Hz) time-courses of neural activity from both the non-BCG corrected (beamforming BCG corrected) and AAS BCG corrected (beamforming+AAS BCG corrected) data using Fieldtrip toolbox (Fieldtrip-20191014, http://www.ru.nl/neuroimaging/fieldtrip) (Oostenveld et al., 2011).

Digitized EEG electrode positions were co-registered with the subjects’ T1-weighted anatomical image using fiducials and scalp surface fitting. A 4-layer (scalp, skull, cerebrospinal fluid (CSF), and brain), anatomically realistic volume conduction BEM head model (Fuchs et al., 2002; Oostendorp and van Oosterom, 1991) was created by segmenting each subject’s T1-weighted images into skin, skull, CSF and brain compartments. In the 4-layer BEM head model, the electrical conductivities of the scalp, skull, CSF and the brain was set to 0.33, 0.0042, 1 and 0.33 S/m, respectively (Gramfort et al., 2010; Kybic et al., 2006). A template grid (5mm spacing) covering the whole brain volume was created from the MNI template brain (ICBM152) and transformed to individual anatomical space of each participant. Potential source locations were then constrained in the gray matter inside each grid. The lead-field matrix using the template grid in individual subject space was then calculated with BEM without constraining the dipole orientations. Then, in order to improve signal to noise ratios (SNRs) of the beamforming spatial filtering, an optimum source orientation/direction was calculated instead of using 3D vector (x, y, z) at the source space location (Sekihara et al., 2004; Sekihara & Nagarajan, 2008). Details of LCMV beamforming spatial filtering technique can be found in the previous articles (Brookes et al., 2008a; 2008b; Uji et al., 2021; van Veen et al., 1997; Westner et al., 2020).

Both non-BCG corrected and AAS BCG corrected data were epoched from −2s to 3s relative to the reviewed spindle onsets for creating beamforming weights and identifying a spindle specific virtual electrode (VE) location. To identify the VE locations of interest related to the spindle neural activity, a pseudo-T statistic was calculated as following, for every location *r* along the 3D grid:

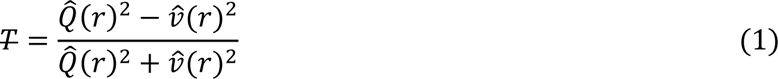

where 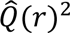 represents the beamformer power estimated in source space location, r, during an active time window, whereas 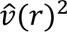 represents the beamformer power estimated in r during a control baseline time window (Brookes et al., 2008a; Hillebrand and Barnes, 2005; Robinson and Vrba, 1999).

Specifically, to detect the optimal spindle VE location, we first calculated broadband (1-100 Hz) time-courses of neural activity during spindle epochs by multiplying the channel level data by the respective broadband beamformer weights in order to attenuate the BCG artefacts through beamforming spatial filtering (Uji et al., 2021). Then the sigma power (11-16Hz) during each spindle duration (group mean spindle duration ± SD = 0.83±0.05s) was extracted, and a corresponding baseline period was defined between −1.5s to −0.5s prior to the spindle onsets. We then identified the maximum peak of the pseudo-T statistic map reflecting the largest sigma band power increase, as the spindle VE location for each subject. From these pseudo-T maps, VE locations were specifically identified for each subject. The mean of the individual subject spindle VE locations during the whole nap was found at [-29 ± 36, 1 ± 33, 40 ± 18] mm [MNI:x,y,z] in the beamforming+AAS BCG corrected data and at [-27 ± 44, 4 ± 27, 44 ± 17] mm [MNI:x,y,z] in the beamforming BCG corrected data (see Fig.1a&b, crosshairs), whereas those during the validated NREM periods was found at [-24 ± 37, −36 ± 30, 62 ± 18] mm [MNI:x,y,z] in the beamforming+AAS BCG corrected data and at [-26 ± 36, −35 ± 30, 56 ± 15] mm [MNI:x,y,z] in the beamforming BCG corrected data (see Fig.1c&d, crosshairs). Since the spindle VE time-course was detected from the central region, the Cz channel signal was recovered for comparison purposes because the Magstim-EGI EEG system used the Cz as a reference electrode for recording.

**Figure 1.**
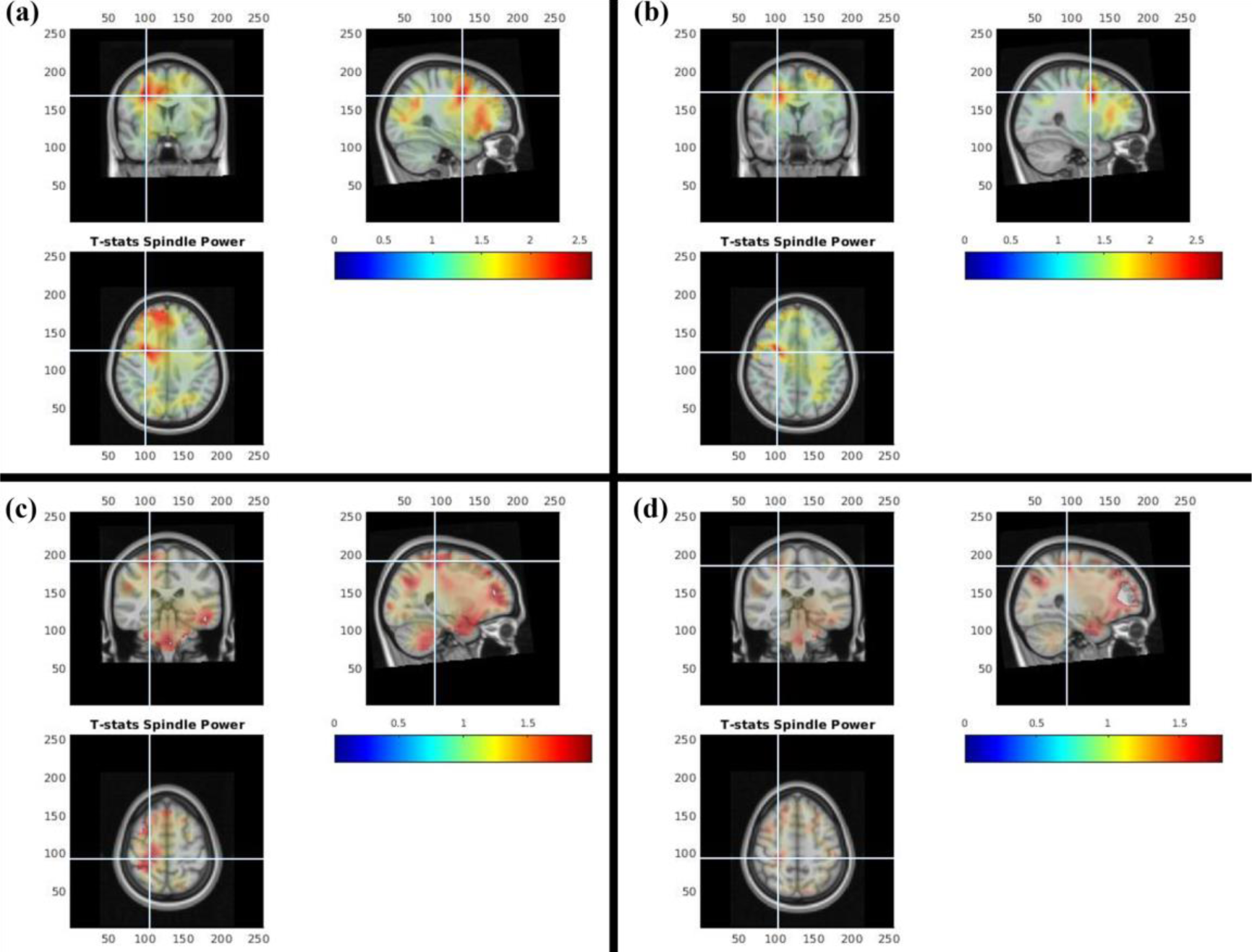
Group average (N = 20) pseudo-T statistic beamformer maps showing regions exhibiting power increases in spindle frequency range (11-16Hz) during the spindle durations (M±SD = 0.83±0.05s) when compared to the baseline window (−1.5 to −0.5 s) in the beamforming+AAS BCG corrected data **(a)** and beamforming BCG corrected data **(b)** during the whole nap, and the beamforming+AAS BCG corrected data **(c)** and beamforming BCG corrected data **(d)** during the validated NREM periods. Each crosshair represents the group average of the individual spindle VE locations **(a)** at [-29 ± 36, 1 ± 33, 40 ± 18] mm [MNI:x,y,z] of the beamforming+AAS BCG corrected date and **(b)** at [-27 ± 44, 4 ± 27, 44 ± 17] mm [MNI:x,y,z] in the beamforming BCG corrected data during the whole nap, and **(c)** at [-24 ± 37, −36 ± 30, 62 ± 18] mm [MNI:x,y,z] in the beamforming+AAS BCG corrected data and **(d)** at [-26 ± 36, −35 ± 30, 56 ± 15] mm [MNI:x,y,z] in the beamforming BCG corrected data during the validated NREM periods.

A broadband (1-100 Hz) time-course of neural activity during the whole MRI scanning period was then calculated from each subject-specific spindle VE locations, using the respective beamforming weights. Specifically, the beamforming weights of the spindle VE were applied to the entire time-course of the whole 187 scalp electrodes during MRI scanning for R-peak locked analysis and spindle locked analysis.

For both R-peak locked analysis and spindle locked analysis, we considered four data sets: non-BCG corrected; AAS BCG corrected; beamforming BCG corrected; beamforming+AAS BCG corrected. These consisted of broadband (1-100 Hz) non-BCG corrected and AAS BCG corrected sensor- and source-level time-courses.

#### R-peak locked analysis

Following the approach we proposed in Uji et al (2021), we first investigated the effects of BCG artefacts on the EEG data quality at the sensor- and source-levels, on both non-BCG corrected and AAS BCG corrected data epoched from −5s to 5s relative to every detected R-peak event (Group mean number (± SD) of R-peak events for the whole nap was 3270.2 (± 849.0) events).

Time-frequency analysis was conducted using a multitaper approach (Scheeringa et al., 2011; Uji et al., 2018) to calculate time-frequency spectrograms of each 10s epoch relative to each individual R-peak event using the Fieldtrip toolbox. Windows of 0.8s duration were moved across the data in steps of 50ms, resulting in a frequency resolution of 1.25 Hz, and the use of three tapers resulted in a spectral smoothing of ±2.5Hz. The calculated time-frequency representations were then transformed in the scale of dB by 10*log10, then demeaned by subtracting the mean power in each frequency during the 10s epoch to calculate power fluctuations around a single R-peak, and averaged across epochs first and then across subjects in both sensor- and source-level.

#### Spindle locked analysis

To investigate the effects of BCG artefacts on the EEG spindle activities at the sensor- and source-levels, both non-BCG corrected and AAS BCG corrected data were epoched from −2s to 3s relative to every reviewed spindle onset (Group mean number (± SD) of spindle events for the whole nap = 160.6 (± 74.6) events & the validated NREM period = 79.2 (± 48.5) events). The group mean spindle density (± SD) was 4.2 (± 2.3) per min during the whole nap.

In the spindle locked time-frequency analysis, time-frequency spectrograms of each 5s epoch relative to spindle events were calculated using the same time-frequency analysis (Scheeringa et al., 2011; Uji et al., 2018). The calculated time-frequency representations were then transformed to dB, then baseline corrected using the baseline period between −1.5s to −0.5s prior to the spindle onsets, and averaged across spindle epochs first and then across subjects in both sensor- and source-level.

To promote new fMRI analysis, we proposed to modulate the detection of every spindle by single-event spindle power values. To do so, separately for each subject, Cz channel and spindle VE time-courses were filtered into the sigma bands (11-16Hz), Hilbert transformed and then the resulting average power during the spindle duration was calculated for each spindle event (Mayhew et al., 2010; Mullinger et al., 2014). These single-event spindle power values represented the trial-by-trial variability in single-event response amplitudes for subsequent GLM analysis of fMRI data.

#### fMRI data analysis

fMRI data were processed using FSL v6.0.5 (https://fsl.fmrib.ox.ac.uk/fsl/). Data from each participant were motion corrected (MCFLIRT), spatially smoothed (5mm FWHM Gaussian kernel), high-pass temporally filtered (100s cut-off), registered to their T1 anatomical brain image (FLIRT), and normalised to the MNI 2mm standard brain. GLM analyses were performed using FEAT v6.00. Based on the spindle events (onset and duration) detected by the trained sleep scorers, first-level analysis was performed employing four different GLM approaches using: 1) the spindle onset and duration (GLM1), 2) spindle onset, duration, and parametric modulation of single-spindle power change estimated from the Cz electrode from the AAS BCG corrected data (GLM2), 3) spindle onset, duration, and parametric modulation of single-spindle power change estimated from the beamforming+AAS BCG corrected data (GLM3) and 4) spindle onset, duration, and parametric modulation of single-spindle power change estimated from the beamforming BCG corrected data (GLM4). All regressors were convolved with the double-gamma HRF. Motion parameters estimated during images realignment (3 translations and 3 rotations around x, y, and z axes) were included in the matrix as confounding regressors of no interest. Additionally, since fMRI BOLD signal amplitudes vary across different vigilance states including sleep (Fukunaga et al., 2006; Fultz et al., 2019; Horovitz et al., 2008; Larson-Prior et al., 2009; Olbrich et al., 2009; Picchioni et al., 2022), the mean fMRI signal inside the white matter regions and the mean fMRI signal in the cerebral spinal fluid regions were included in the GLM model as nuisance regressors (Caballero-Gaudes and Reynolds, 2017; Fang et al., 2020). These first-level results were then combined and analysed across all subjects at the group-level using FSL randomize non-parametric permutation testing with 5,000 permutations (Eklund et al., 2016; Winkler et al., 2014). The resulting group-level BOLD Z statistic images were thresholded using Gaussian random field theory based maximum height thresholding (Worsley, 2012) with the voxel-wise inference (p < 0.05 corrected). These four proposed fMRI analyses were applied to both fMRI data during the whole nap and validated NREM periods.

### Data and code availability

All data used in this study will be available via a request to the corresponding author (M.U.). Open access software used in this study is available at https://sccn.ucsd.edu/eeglab (EEGLAB), http://www.ru.nl/neuroimaging/fieldtrip (FieldTrip), and https://fsl.fmrib.ox.ac.uk/fsl (FSL). Custom analysis code used in this study will be available via a request to the corresponding author (M.U.).

## Results

### Sleep architecture & sleep spindle characteristics

As shown in Table 1, all 20 participants experienced NREM1 and NREM2 sleep, 16 had NREM3, and no one experienced REM sleep period during the recovery nap. The group mean total sleep duration was 46.60 (SD = 14.48) min during the 60min recovery nap in the MRI scanner. The average sleep latency of these 20 participants was 5.32 ± 9.77 min. Sleep fragmentation was also calculated as the total number of awakenings and shifts to NREM1 divided by the total sleep time (sleep fragmentation index: SFI). The group mean SFI was 8.91 events per hour (SD = 5.62). Figure S3 revealed each subject’s sleep hypnogram during the whole nap EEG-fMRI session. The participants had on average 168.65 sleep spindles (SD = 70.10, min = 51, max = 293) during NREM sleep of the 60min nap. The group mean spindle density was 4.23 (SD = 2.34) per minute, and the group mean spindle duration was 0.83s (SD = 0.05). For the validated NREM periods (5 min selected for each subject), the participants had on average 79.20 (SD = 48.54, min = 7, max = 159) sleep spindles, and the group mean spindle duration during the validated NREM periods was 0.84s (SD = 0.06). The group mean spindle density during the validate NREM period was 7.91 (SD = 4.35) per minute.

### BCG artefact corrections

#### R-peak locked analysis

We detected on average 3270.2 (SD=849.0) R-peak events during the whole nap recording. Figure 2a illustrates the group mean time-frequency representations (TFRs) time-locked at the individual R-peak events for non-BCG corrected and AAS BCG corrected channel (Cz) signals, as well as beamforming BCG corrected and beamforming+AAS BCG corrected source signals estimated on spindle VE. When considering non-BCG corrected data at Cz, the power fluctuations of the BCG artefact were observed at the peak power of around +2dB, especially below 20Hz, at the time of R-peaks. After applying the standard AAS BCG correction, these power fluctuations were reduced to the peak power of around 0.7dB at the time of R-peaks especially below 20Hz. Furthermore, these BCG artefacts are reduced to 1dB with the beamforming BCG correction and 0.3dB with the beamforming+AAS BCG correction. These effects can also be observed in the power spectrum calculated for the periods both between 0 to 0.5s and between 0.5 to 1s after the individual R-peak events (Fig.2b). The BCG denoising performance of the beamforming+AAS correction was superior among the other methods, especially for frequencies below 20Hz, therefore particularly important to enhance the spindle frequency band.

**Figure 2.**
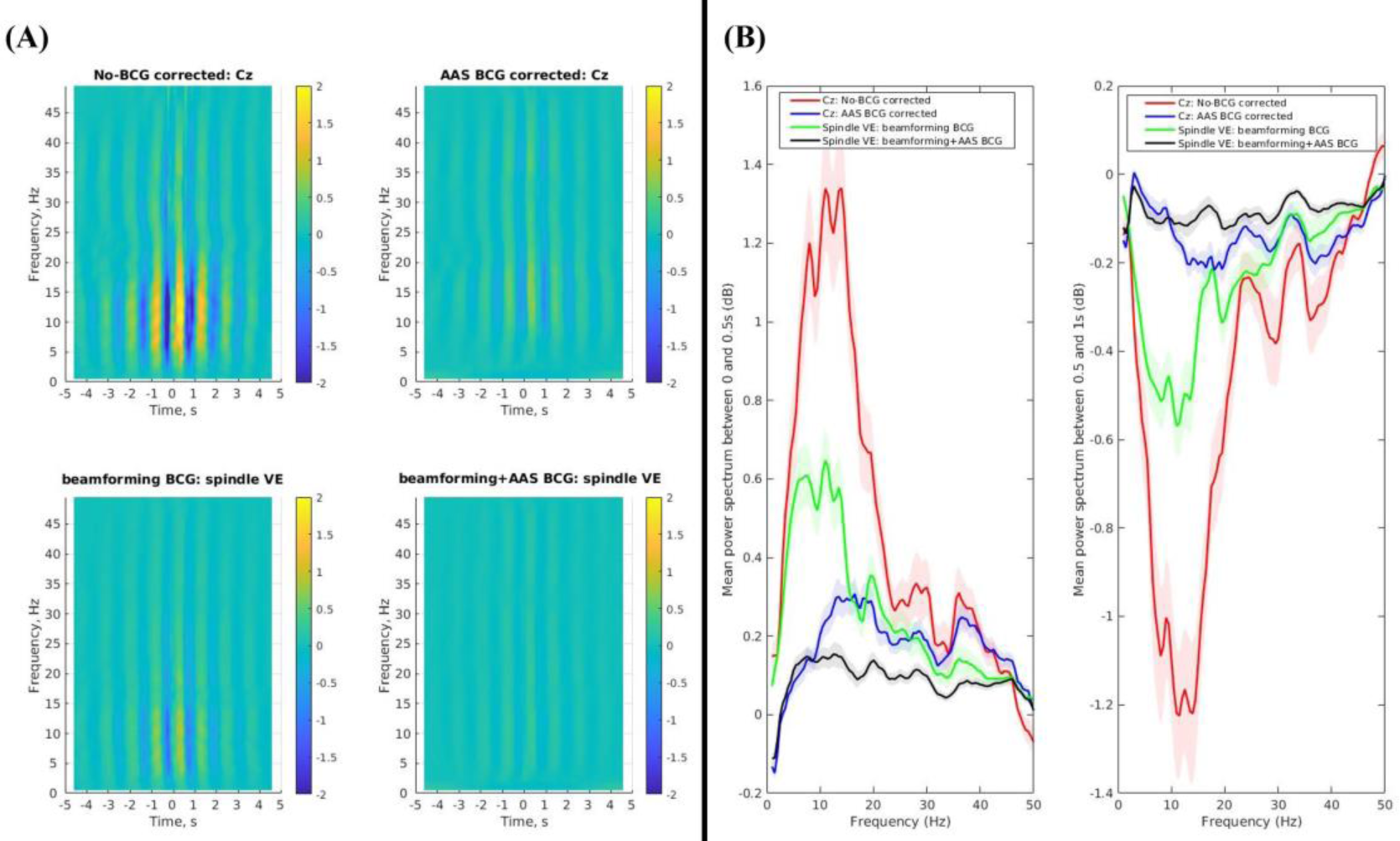
**(A)** Group mean (N=20) time-frequency representations (TFRs) of R-peak locked epochs during the whole nap for the non-BCG corrected (Cz: top left), AAS BCG corrected (Cz: top right), beamforming BCG corrected (spindle VE: bottom left), and beamforming+AAS BCG corrected data (spindle VE: bottom right). These TFRs demonstrated −5s to 5s relative to the R-peak event onsets (t=0). **(B)** Group mean (N=20) power spectrums during the time-period of 0-0.5s (left) and the time-period of 0.5-1s (right) relative to the R-peak event onsets during the whole nap.

#### Spindle locked analysis

Figure 3a shows the group mean TFRs time-locked to the individual spindle events during the whole nap for non-BCG corrected and AAS BCG corrected channel (Cz) signals, and beamforming BCG corrected and beamforming+AAS BCG corrected source signals estimated on spindle VE. When considering non-BCG corrected data at Cz, apparent sigma (11-16Hz) power increase could not be observed during the spindle durations (group mean = 0.83±0.05s). However, after applying the standard AAS BCG correction, the sigma power increase was evident at Cz. Furthermore, both beamforming BCG correction and beamforming+AAS BCG correction also successfully recovered the spindle activity from the BCG artefacts. These findings can be confirmed in the power spectrum when each participant’s spindle power spectral density was calculated for the spindle durations for each spindle, and then averaged across the number of spindles (Fig.3b). The group mean spindle power (11-16Hz) was −0.29dB in the non-BCG corrected Cz, 0.78dB in the AAS BCG corrected Cz, 0.50dB in the beamforming BCG corrected spindle VE, and 0.64dB in the beamforming+AAS BCG corrected spindle VE.

**Figure 3.**
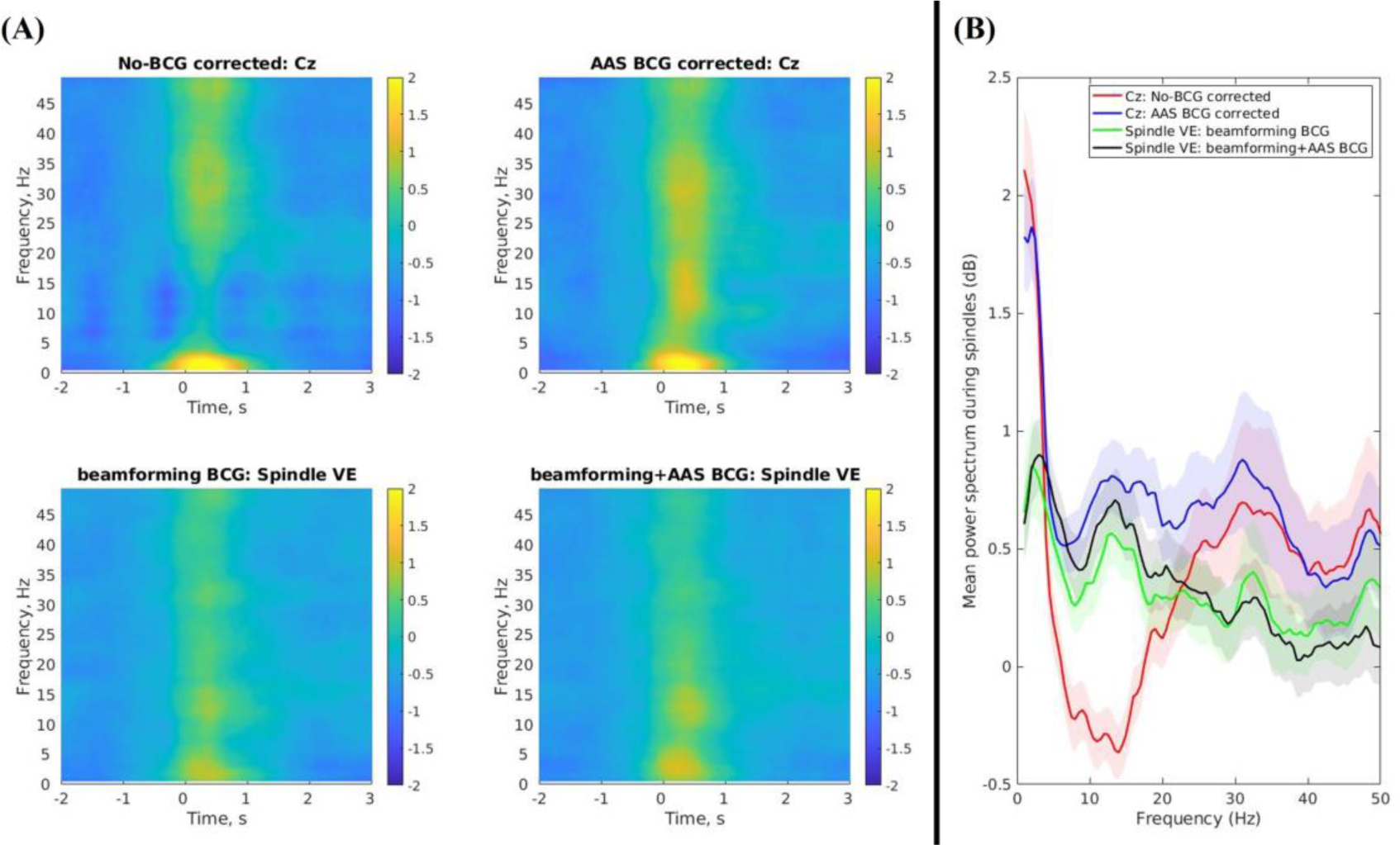
**(A)** Group mean (N=20) time-frequency representations (TFRs) of spindle activity during the whole nap for the non-BCG corrected (Cz: top left), AAS BCG corrected (Cz: top right), beamforming BCG corrected (spindle VE: bottom left), and beamforming+AAS BCG corrected data (spindle VE: bottom right). These TFRs demonstrated −2s to 3s relative to the spindle onsets (t=0). **(B)** Group mean (N=20) power spectrums of the detected spindle durations during the whole nap (M±SD = 0.83±0.05s).

We then investigated the validated spindles in the clear NREM (N2 & N3) periods, where the four trained scorers were allowed to add and remove the spindle events if the automatic spindle detection failed, in addition to accepting or rejecting the automatically detected spindles during the other periods. During the validated NREM periods, we detected 79.2 (SD = 48.5) spindle events with the group mean duration of 0.84s (SD=0.06) and the group mean spindle density of 7.91 (SD = 4.35) per minute. Supplementary Figure S3a shows the group mean TFRs time-locked at the individual spindle events during the validated NREM periods. When considering non-BCG corrected data at Cz, again the apparent sigma (11-16Hz) power increase was not evident during the spindle durations (group mean = 0.84±0.06s). However, after applying the standard AAS BCG correction, the sigma power increase can be observed at Cz. Furthermore, both beamforming BCG correction and beamforming+AAS BCG correction successfully recovered the spindle activity from the BCG artefacts. These findings can also be confirmed by the power spectrum calculated for the spindle durations (Supplementary Fig.3b). Specifically, the group mean spindle (11-16Hz) power was −0.30dB in the non-BCG corrected Cz, 0.96dB in the AAS BCG corrected Cz, 0.37dB in the beamforming BCG corrected spindle VE, and 0.51dB in the beamforming+AAS BCG corrected spindle VE.

Together, these findings support that the spindle activity during EEG-fMRI were heavily influenced by the presence of the BCG artefacts, and the EEG signals need to be denoised carefully. Furthermore, all BCG correction methods successfully preserved the spindle activity from the BCG artefacts.

### fMRI GLM analysis

Since adding single-spindle event variability into the GLM model was expected to improve sensitivity and specificity of the fMRI GLM analysis, we performed four different GLM analyses based on the spindle events (onset and duration) reviewed by the trained sleep scorers, using fMRI data during the whole nap. The GLM included: 1) the spindle onset and duration (GLM1), 2) spindle onset, duration, and parametric modulation of single-spindle power change estimated from the Cz electrode from the AAS BCG corrected data (GLM2), 3) spindle onset, duration, and parametric modulation of single-spindle power change estimated from the beamforming+AAS BCG corrected (GLM3) and 4) spindle onset, duration, and parametric modulation of single-spindle power change estimated from beamforming BCG corrected data (GLM4). As expected across 20 subjects, we observed a significant main-effect positive BOLD response to the spindle activities (voxel-wise corrected p<0.05) for all four GLM models in the thalamus, caudate, posterior cingulate, hippocampus, temporal cortex, cerebellum, and brainstem (GLM1 in Fig.4, GLM2 in Fig.5, GLM3 in Fig.6, GLM4 in Fig.7). In addition, the BOLD responses in the thalamus, caudate, posterior cingulate, hippocampus, temporal cortex, cerebellum, and brainstem demonstrated a significant positive correlation with single-event spindle power change (voxel-wise corrected p<0.05) at the Cz (GLM2) and spindle VEs (GLM3, GLM4).

**Figure 4.**
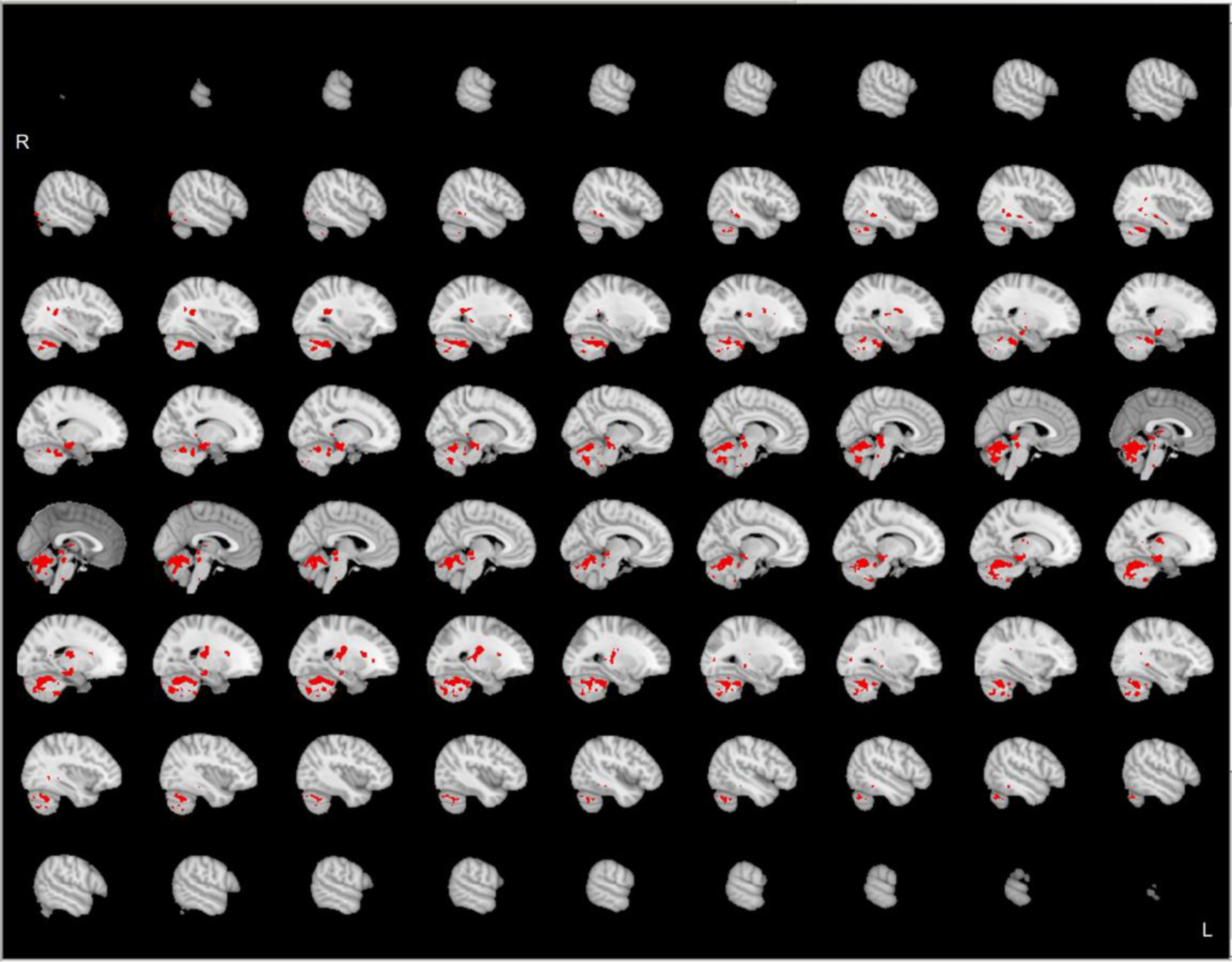
Group-level (N=20) significant main effects for GLM1 (spindle onset & duration) during the whole nap, revealed by FSL randomize non-parametric permutation testing with 5,000 permutations (Eklund et al., 2016; Winkler et al., 2014) with the voxel-wise inference (p < 0.05 corrected).

**Figure 5.**
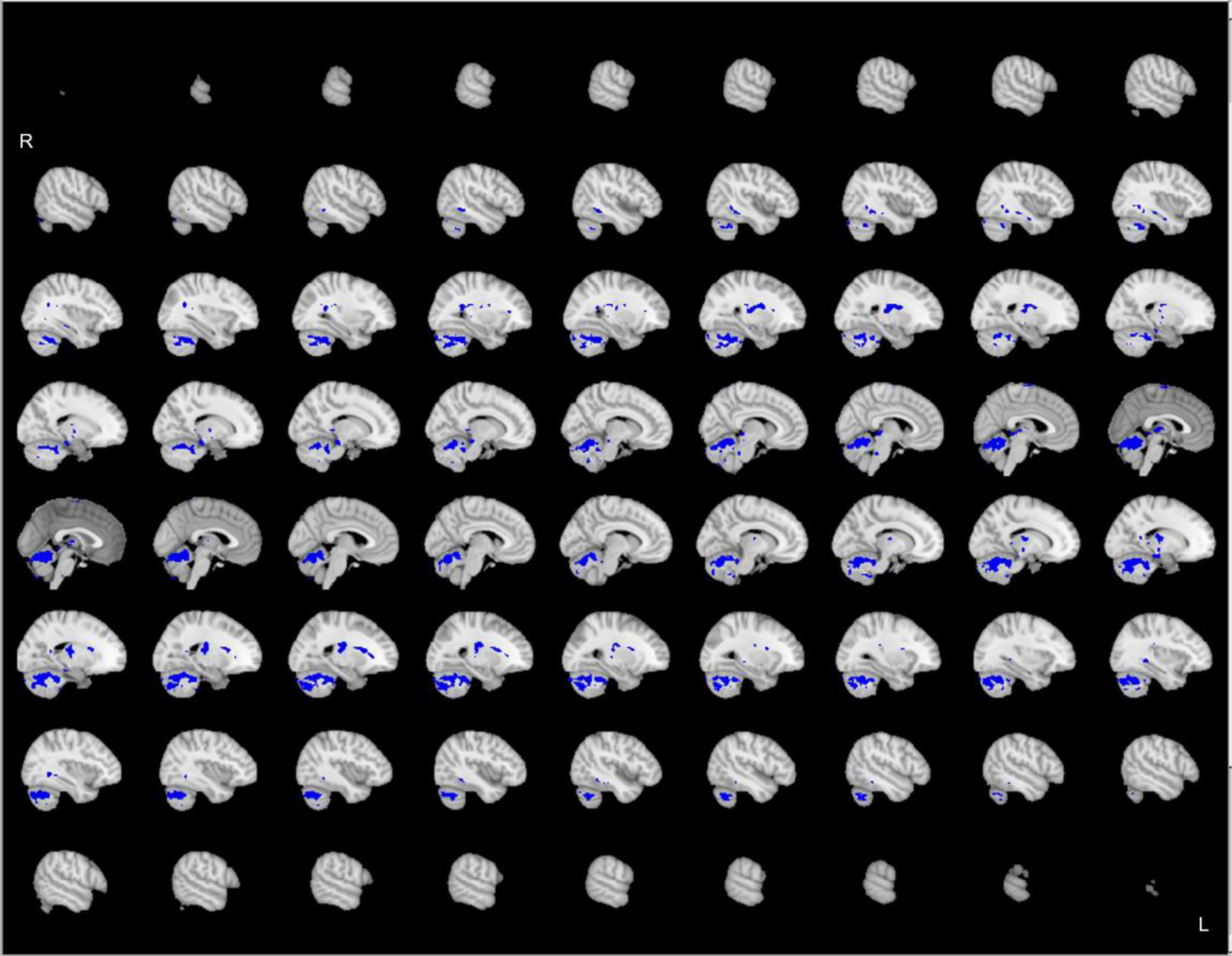
Group-level (N=20) significant main effects for GLM2 (spindle onset, duration, & single-spindle power changes from the Cz electrode of the AAS BCG corrected data) during the whole nap, revealed by FSL randomize non-parametric permutation testing with 5,000 permutations (Eklund et al., 2016; Winkler et al., 2014) with the voxel-wise inference (p < 0.05 corrected).

**Figure 6.**
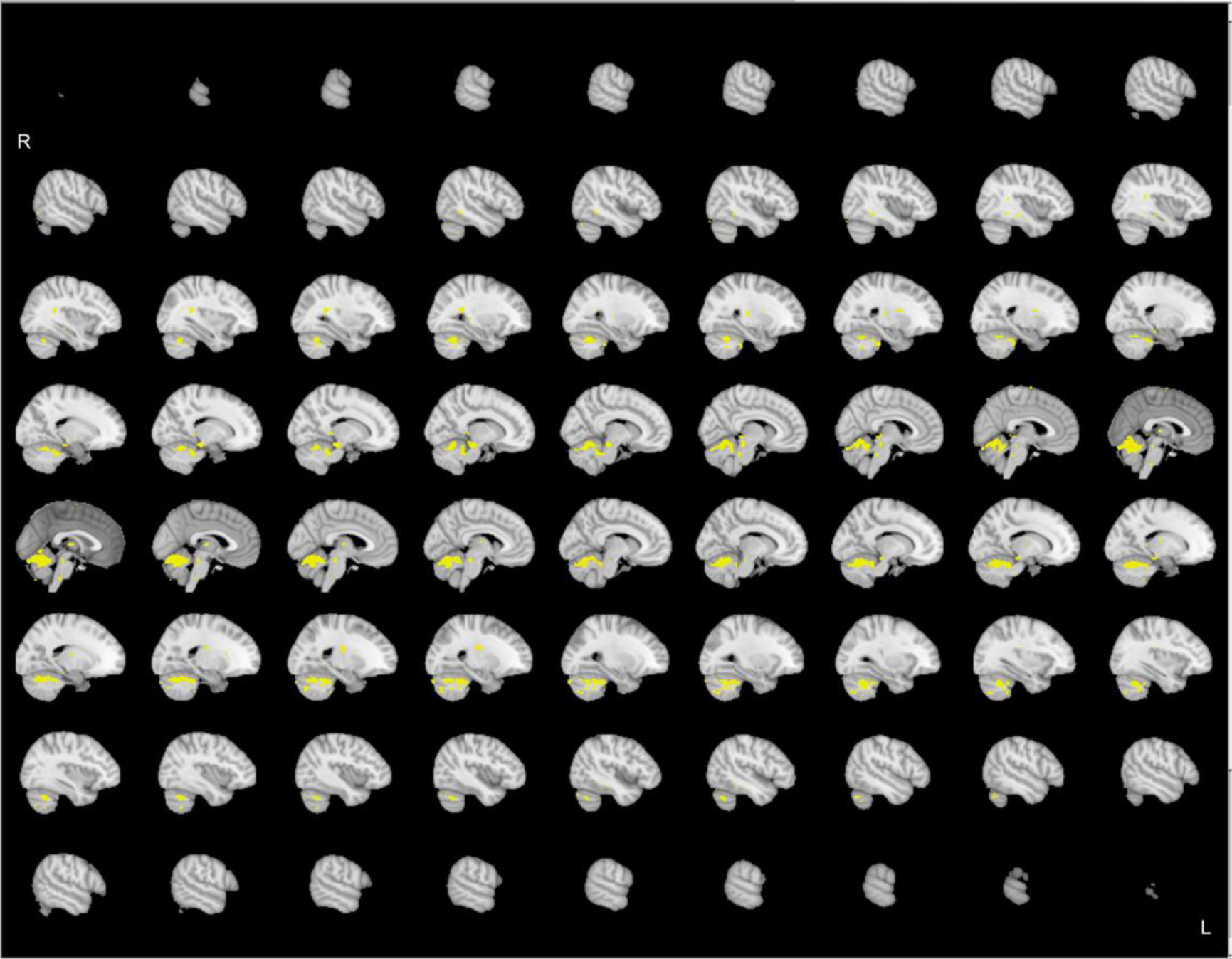
Group-level (N=20) significant main effects for GLM3 (spindle onset, duration, & single-spindle power changes from the spindle VE of the beamforming+AAS BCG corrected data) during the whole nap, revealed by FSL randomize non-parametric permutation testing with 5,000 permutations (Eklund et al., 2016; Winkler et al., 2014) with the voxel-wise inference (p < 0.05 corrected).

**Figure 7.**
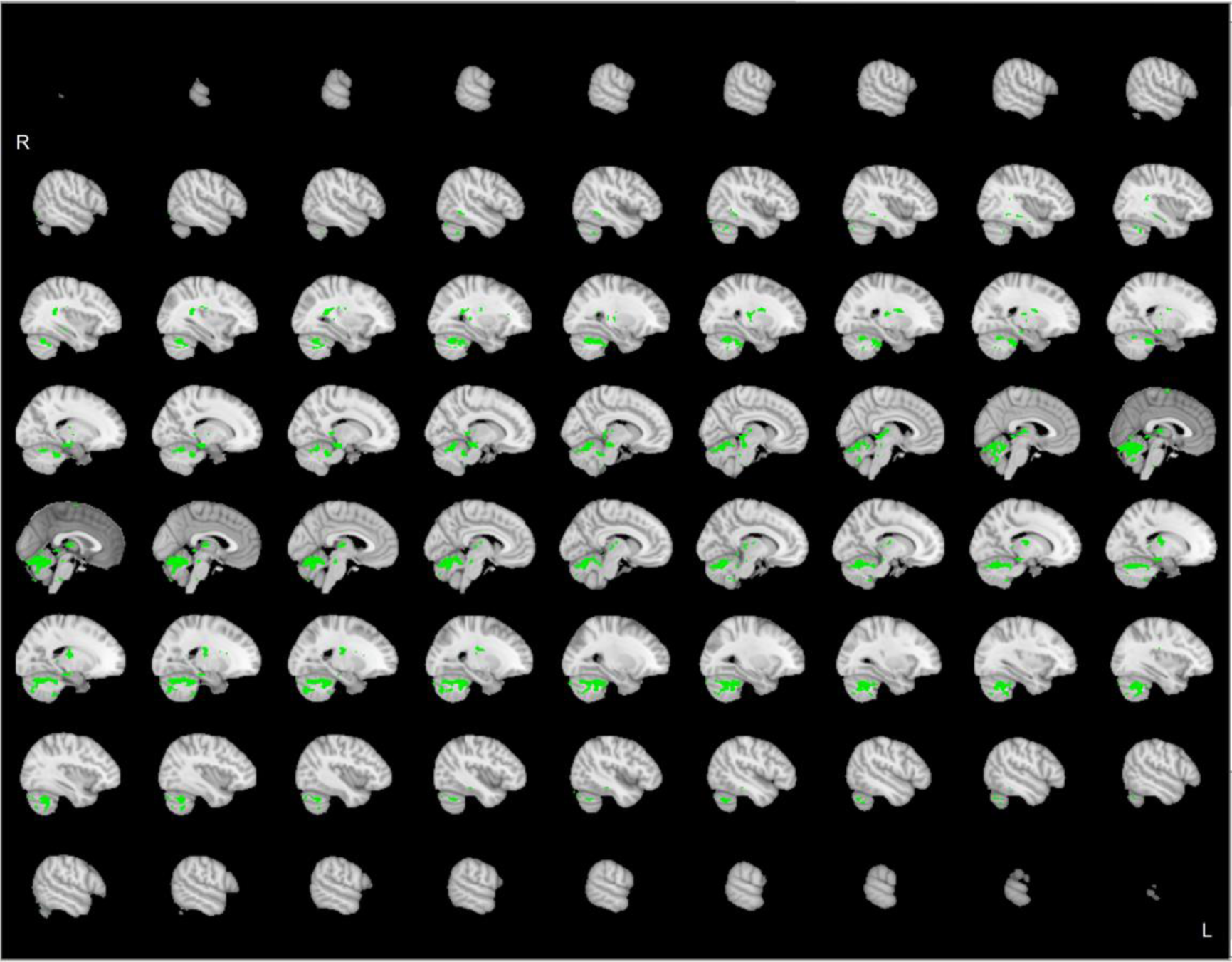
Group-level (N=20) significant main effects for GLM4 (spindle onset, duration, & single-spindle power changes from the spindle VE of the beamforming BCG corrected data) during the whole nap, revealed by FSL randomize non-parametric permutation testing with 5,000 permutations (Eklund et al., 2016; Winkler et al., 2014) with the voxel-wise inference (p < 0.05 corrected).

The total number of significant clusters (p<0.05 corrected) was 55 in GLM1, 54 in GLM2, 50 in GLM3, and 48 in GLM4, whereas the total number of significant gray matter voxels/the total number of all voxels was 1595/4984 in GLM1, 1582/5542 in GLM2, 1103/3316 in GLM3, and 1395/4501 in GLM4. Furthermore, the number of the overlapping regions among the four significant maps was 1977 voxels, and the percentage of the overlapping regions over the total number of significant voxels was 39.7% in GLM1, 35.7% in GLM2, 59.6% in GLM3, and 43.9% in GLM4 (Fig.8). The GLM3 provided the most intersected regions during the whole nap among the four different GLM models.

**Figure 8.**
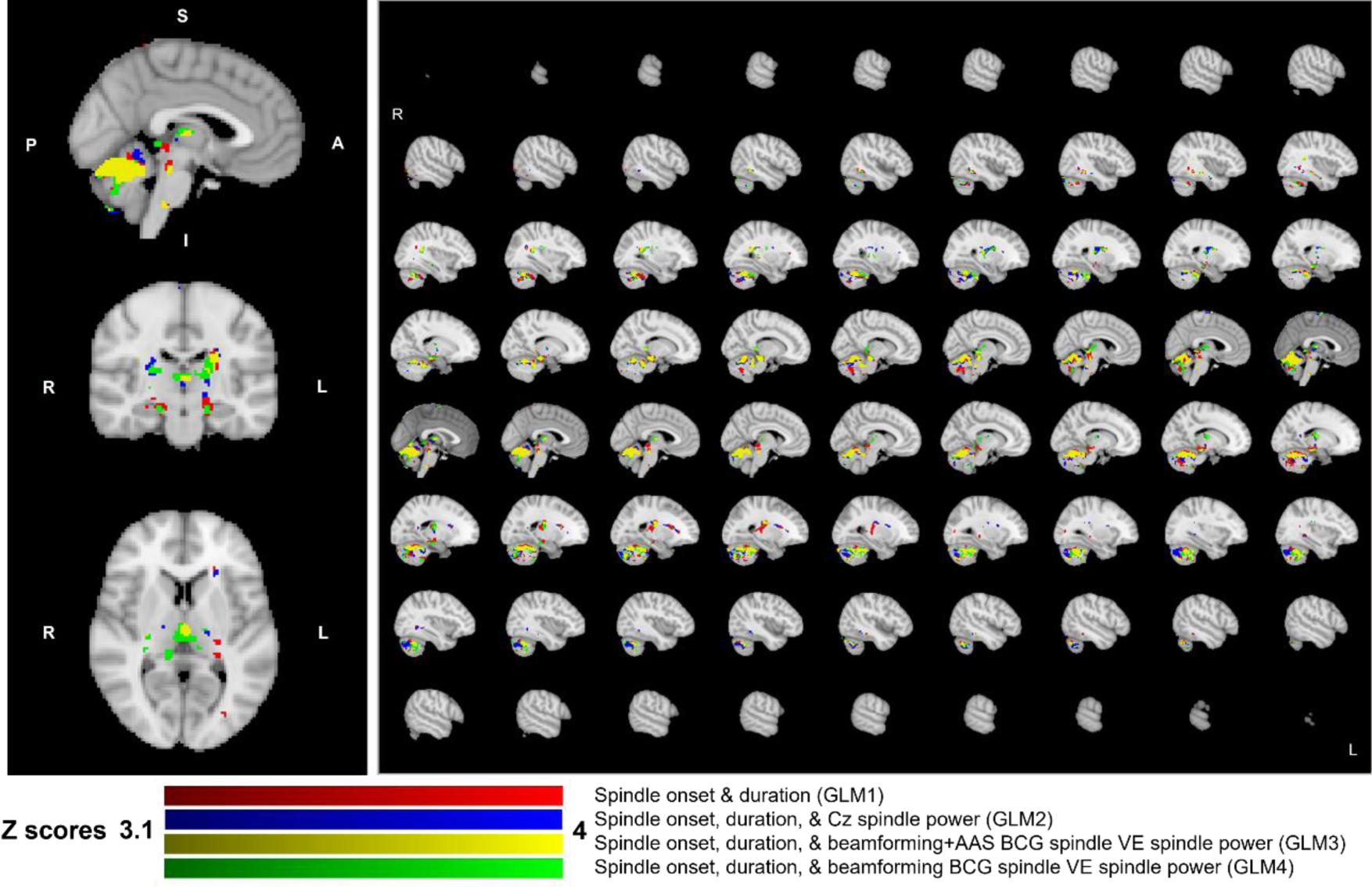
Overlapping brain regions between four different GLM models during the whole nap: GLM1 (spindle onset & duration), GLM2 (spindle onset, duration, & single-spindle power changes from the Cz electrode of the AAS BCG corrected data), GLM3 (spindle onset, duration, & single-spindle power changes from the spindle VE of the beamforming+AAS BCG corrected data), GLM4 (spindle onset, duration, & single-spindle power changes from the spindle VE of the beamforming BCG corrected data). The GLM3 provided the most concordant regions during the whole nap among the four different GLM models.

Similarly, during the validated NREM periods, even if shorter data were considered for each participant (5 min sections), we still found a significant main-effect positive BOLD response to the spindle activities (voxel-wise corrected p<0.05) for all four GLM models in the thalamus and cerebellum (GLM1 in Fig.S5, GLM2 in Fig.S6, GLM3 in Fig.S7, GLM4 in Fig.S8). Although the validated NREM periods revealed only the thalamus and cerebellum regions, the BOLD responses were positively correlated with single-event spindle power fluctuations (voxel-wise corrected p<0.05) at the Cz (GLM2) and spindle VEs (GLM3, GLM4) in the similar manner as the fMRI GLM analyses for the whole nap.

## Discussion

In this study, we first investigated the effect of the beamforming technique to attenuate ballistocardiogram (BCG) artefacts in EEG-fMRI data acquired during sleep, even without detecting cardiac pulse (R-peak) events from simultaneous ECG recordings. In a second step, we quantified how this technique would reconstruct sleep spindle activity (11-16Hz) normally occurring in NREM sleep, which would be heavily affected by the BCG artefacts. Including our previous work (Uji et al., 2021), several studies have demonstrated that beamformer is highly efficient in attenuating artefactual signals which have different spatial origins from the underlying signal of interest such as eye movements (Cheyne et al., 2006), orthodontic metal braces (Cheyne et al., 2007) in MEG, and the MRI-related artefacts (i.e., BCG artefacts) (Brookes et al., 2009, 2008a; Mullinger and Bowtell, 2011; Uji et al., 2021). Specifically, the beamforming spatial filter rejects sources of signal variance that are not concordant in source locations inside the brain based on the forward model, and attenuates all unwanted source activities outside of the specified source location of interest in the data (e.g., eye movements, BCG) without having to specify the location or the configuration of these unwanted underlying source signals (Brookes et al., 2007; Huang et al., 2004; van Veen et al., 1997). Since the beamforming spatial filtering appears as a promising denoising technique in the context of EEG-fMRI studies (Brookes et al., 2009, 2008a; Mullinger and Bowtell, 2011), we hypothesised that the beamforming technique would attenuate the BCG artefacts of the EEG-fMRI acquired during sleep to a similar extent as compared to the conventional AAS BCG artefact corrections. Moreover, we expected that the beamforming approach would accurately estimate single-event spindle activity while supressing the BCG artefacts when compared to channel level signals, and further improve the sensitivity and specificity of the fMRI GLM analysis when modelling the single-event spindle activity variability in the fMRI GLM regressors (Mullinger et al., 2017, 2014, 2013; Pakenham et al., 2020; Uji et al., 2018; Wilson et al., 2019).

### BCG corrections in EEG-fMRI acquired during sleep

In the time-frequency analysis of the R-peak locked events (Figure 2), we first showed that generally the standard AAS and beamforming related methods (beamforming BCG & beamforming+AAS BCG correction) suppressed BCG artefacts as compared to non-BCG corrected data. Clearly, the BCG artefacts and its residuals obscured EEG signals lower than 20Hz (Figure 2). If the BCG corrections are suboptimal, this would become more problematic for sleep spindle (11-16Hz) detections and analysis in the EEG-fMRI data.

We then evaluated the impact of beamforming spatial filtering to recover spindle activity, when applied to AAS BCG corrected (beamforming+AAS BCG corrected) and non-BCG corrected (beamforming BCG corrected) data, by comparing them with sensor level data (AAS BCG corrected & non-BCG corrected data). The individual virtual electrode (VE) locations were determined by the maximum increase in spindle power (11-16Hz) in the source space (spindle VE) optimizing the sigma power increase during the spindle duration as compared to the baseline. We then obtained the location of these spindle VEs for each subject and found an overall concordance across subjects (Fig.1). The location of the spindle VEs was consistent to the ones reported in the literature (Brancaccio et al., 2020; Klinzing et al., 2016; Manshanden et al., 2002; Zerouali et al., 2014). We demonstrated that the spindle VEs time-courses from both beamforming approaches (beamforming BCG corrected; beamforming+AAS BCG corrected) contained less BCG artefacts than the uncorrected data at the sensor level, preserving the spindle activity by suppressing the BCG artefact (Fig.2&3).

Overall, our findings demonstrated a consistent denoising performance of the beamforming BCG corrections (both beamforming BCG & beamforming+AAS BCG correction) when considering sleep data, similarly to EEG correction during task data as previously reported (Brookes et al., 2008a; Uji et al., 2021). Our results support and extend on the previous findings related to sleep and longer duration EEG-fMRI acquisitions (up to 60mins for this dataset). When a long recording is required during sleep, R-peak event detection based on visual inspections might become time-consuming and more problematic due to potential increases in ECG signal distortion and saturation. Moreover, heart rates also vary across vigilance levels during sleep (Tobaldini et al., 2013), resulting in more dynamical change of the cardiac responses during sleep when compared to the awake or task conditions. In such cases, AAS approach might not be an optimal solution for the BCG correction. Our findings revealed that our proposed data-driven beamforming denoising approach, which does not require the detection of R-peak events from ECG recording, is promising in attenuating the BCG artefacts in the EEG-fMRI acquired during sleep, while preserving important sleep-related EEG signatures such as sleep spindles.

### fMRI GLM analysis

One advantage of applying the proposed beamforming technique in EEG-fMRI is to improve the sensitivity and specificity of fMRI BOLD general linear model (GLM) analysis for sleep studies. Motivated by the previous findings integrating the single-event EEG variability in the fMRI GLM analysis in cognitive and sensory tasks (Mullinger et al., 2017, 2014, 2013; Pakenham et al., 2020; Uji et al., 2018; Wilson et al., 2019) and in epilepsy (Grouiller et al., 2011; LeVan et al., 2010; Vulliemoz et al., 2010), we investigated whether here proposed beamforming technique could benefit the fMRI GLM analysis. Specifically, this can be achieved by extracting a source activity from a selected VE location, calculating the Hilbert transformed envelope of source neural activities in a specific frequency band, and parameterizing regressors with the Hilbert envelope amplitude fluctuations instead of using consistent fixed binary boxcar (i.e., 0 or 1) for the fMRI GLM analysis (Mullinger et al., 2014; Uji et al., 2018). This is based on the findings demonstrating that the amplitudes and variabilities of BOLD responses have been better explained by this parameterized GLM (i.e., single-trial/event correlation analysis), doing so by better taking into account the source of neural variability in EEG responses, than by a conventional GLM of consistent amplitude responses (Mullinger et al., 2014; Vulliemoz et al., 2010). This Hilbert transformed envelope has been used in the different frequency bands (alpha 8–13 Hz: Mullinger et al., 2013, 2014; Mullinger, Cherukara, Buxton, Francis, & Mayhew, 2017; beta 13– 30 Hz: Pakenham et al., 2020; Wilson et al., 2019; gamma 55–80 Hz: Uji et al., 2018). For example, this parameterized GLM analysis improved the specificity and sensitivity to identify the neurophysiological origin of the negative BOLD response to unilateral median nerve stimulation (Mullinger et al., 2014) and negative BOLD responses between stimulated and unstimulated sensory cortices (Wilson et al., 2019). Furthermore, the parameterized GLM analysis revealed a positive correlation between single trial gamma band variability and BOLD responses over the contralateral primary motor cortex during finger abduction movements, indicating a tight neurovascular coupling between the gamma band activity and BOLD responses (Uji et al., 2018). Here, we hypothesized that such parameterized fMRI GLM approach could be used to better localize the neurovascular coupling of sleep specific neural activities (i.e., spindle, slow-wave) in sleep research, after the data-driven beamforming denoising technique suppresses the BCG artefacts which remain problematic for the frequency of sleep specific neural activities.

In the present EEG-fMRI data acquired from 20 healthy subjects, we estimated single-event spindle power changes (11-16Hz) both from the channel- (Cz) and source-level (spindle VE) and investigated to which degree the spindle-related BOLD signal amplitude change and variability could be explained by such power changes. Especially, we assessed four different GLM models (GLM1-4). Using all four GLM models, we observed the main effect includes a significant positive BOLD response to the spindle activity in the thalamus, caudate, posterior cingulate, hippocampus, temporal cortex, cerebellum, and brainstem (Fig.4, Fig.5, Fig.6, Fig.7). These brain regions were indeed expected to be recruited during spindles and consistent with previous findings (Dang-Vu et al., 2011; Fang et al., 2019; Jegou et al., 2019; Schabus et al., 2007). Since EEG or MEG source imaging still finds it difficult to accurately localize the deep structure activities (i.e., thalamus, hippocampus, brainstem, caudate), simultaneous EEG-fMRI recording is crucial to advance our understanding of the subcortical brain activities in humans.

In addition, if the beamforming approach would allow us to estimate the single-event spindle power changes more accurately in the source level when compared to the channel level, we expected the GLM3 and GLM4 would improve the specificity of the fMRI GLM analysis when compared to GLM1 and GLM2. We especially expected that this beamforming approach would better localize the recruited regions as demonstrated for the previous EEG-fMRI studies in focal epilepsy (Grouiller et al., 2011; LeVan et al., 2010; Vulliemoz et al., 2010). Even if we could not have access to an accurate ground truth to evaluate these fMRI results, we nevertheless found the GLM3 and GLM4 exhibited fewer number of significant voxels when compared to the GLM1 and GLM2. Specifically, the number of significant voxels outside the gray matter regions in the GLM3 and GLM4 were fewer than the GLM1 and GLM2, suggesting an improvement in the specificity to localize the origins of neurovascular coupling related with spindle activity. Interestingly, the GLM2 (spindle onset, duration, & single-spindle power changes from the Cz of the AAS BCG corrected data) increased the number of significant voxels both in total and outside of the gray matter regions without any change inside the gray matter when compared to the GLM1. Specifically, adding the single-event spindle power change in the channel level might have hindered the specificity of the fMRI GLM analysis. This might be due to less accurate estimations of each spindle power change due to the residual BCG artefacts, which can also be confirmed when extracting spindle event power amplitude variability (Fig.S9). In Supplementary Figure S9, we illustrated all spindle power amplitudes across 20 subjects during the whole nap and the validated NREM periods in the histogram. 95% confidence intervals of the channel level spindle power amplitude (AAS BCG corrected) were ±19 during the whole nap and ±4 during the validated NREM periods, whereas those in both source levels (beamforming BCG corrected; beamforming+AAS BCG corrected) were ±4 during the whole nap and ±1 during the validated NREM periods for both source level corrections. These larger amplitude changes at the channel-level might be because the residual BCG artefacts might have caused overestimation of each spindle amplitude resulting in less accurate estimates for the GLM regressors as compared to the source level estimates (GLM3 & GLM4) or just a boxcar ([1 0], GLM1) regressor.

### Benefit of beamforming approach to EEG-fMRI acquired during sleep

Although several EEG source imaging techniques have been used in the EEG-fMRI during cognitive and sensory tasks (LORETA: Whittingstall et al., 2010; beamforming: Brookes et al., 2009, 2008a; Mullinger et al., 2017, 2014; Uji et al., 2018; Wilson et al., 2019), in epilepsy research (LAURA: Groening et al., 2009; Vulliemoz et al., 2010, 2009), and in resting-state research (Minimum Norm Estimate (MNE): Wirsich et al., 2021), these techniques have been rarely applied to EEG data recorded in the MRI scanner during sleep. In our knowledge, only one previous study (Boutin et al., 2018) used depth-weighted minimum norm estimate (wMNE) in order to investigate the functional role of NREM2 sleep spindles in motor skill consolidation after motor sequence learning. This lack of attempts applying source imaging to EEG-fMRI in general, more especially in sleep research, could be associated with the lack of ability to properly correct MRI-related artefacts. Even after careful artifact corrections using different approaches (Allen et al., 1998, 2000; Iannotti et al., 2015; Mullinger et al., 2008b; Vanderperren et al., 2010), EEG data inside the MRI scanner remains with lower signal quality when compared to EEG data recorded outside the scanner. In general, EEG source imaging procedures require good EEG quality over the whole spatial topography (Leijten and Huiskamp, 2008). Furthermore, most studies aimed at detecting sleep related EEG features to be analysed with the fMRI framework (e.g., spindles, slow waves) using only low-density EEG layout (less than 32 channels), therefore limiting the possibility to perform reliable and accurate EEG source reconstructions. To the best of our knowledge, our study is the first one to investigate the benefit of using beamforming approach in EEG-fMRI acquired during sleep, while benefitting from high-density EEG recordings. In this study, we demonstrated that applying beamforming spatial filtering technique reduced the BCG artefacts by half and successfully recovered the spindle activity even without relying on R-peak detections (Fig.2&3). A commonly used BCG artefact correction method is average artefacts subtraction (AAS) requiring high precisions of cardiac pulse (R-peak) event detections from simultaneous ECG recording which are used for subtracting averaged BCG artefact templates (Abreu et al., 2018; Allen et al., 2000, 1998; Bullock et al., 2021). However, the ECG signal in the MRI scanner is also often distorted, which makes automatic detection of R-peaks problematic (Chia et al., 2000; Iannotti et al., 2015; Mullinger et al., 2008b), and requires manual correction which is significantly time-consuming for a long recording such as sleep (i.e., maximum up to 2 hours). Additionally, heart rates during sleep recording will be varied across different vigilant stages (Tobaldini et al., 2013). Thus, detecting R-peak events from the ECG is difficult, so that this procedure may sometimes become unreliable.

Findings reported from our R-peak and spindle locked analyses demonstrated that our both proposed beamforming approaches (beamforming+AAS BCG corrected & beamforming BCG corrected data) successfully recovered the detected spindle activity during the whole nap and validated NREM periods, while minimizing the effects of the BCG artefacts. After suppressing the BCG artefacts, other residual artefacts (around 30Hz) also appear to be minimized through both beamforming approaches when compared to standard AAS BCG correction. This would help detecting spindle activity more efficiently and in a robust manner, when compared to background activity due to enhancing the signal to noise ratios of the spindle events. Indeed, in a recent MEG study, Hill et al. (2020) reported similar beamforming source reconstruction approach improving the SNRs by over 1.5 times for the motor beta frequency activity during finger abduction movements and for the visual gamma (55-70 Hz) frequency activity during the visual stimuli when compared to those in the sensor space. In our previous study (Uji et al., 2021), we also reported the improvement of SNR in EEG data (i.e., alpha activity in visual cortex & beta activity in motor cortex) by minimizing any other residual artefacts caused by fMRI acquisition. These findings are consistent with the fact that beamforming source reconstruction will improve the SNRs when compared to the sensor level (Brookes et al., 2009; Hill et al., 2020; Sekihara et al., 2004), and this approach will be beneficial especially for EEG–fMRI analyses (Brookes et al., 2009; Mullinger & Bowtell, 2011).

Moreover, in agreement with the previous work (Brookes et al., 2008a; Uji et al., 2021), an optimal strategy to suppress the BCG artefacts, while preserving brain signals for the spindle-locked analysis and fMRI GLM analysis, was to apply the beamforming approach after the AAS BCG correction (beamforming+AAS BCG correction) as a post-processing procedure. This could be because the large part of the BCG artefacts is removed by the AAS approach if the R-peak detections are precise and accurate and then, the beamforming further attenuates the residual BCG artefacts as suggested by Mullinger and Bowtell (2011). It is worth mentioning that hardware-based solutions (e.g., Reference layer artefact subtraction: Chowdhury et al., 2014; Carbon-wire loop: Masterton et al., 2007; van der Meer et al., 2016; Optical Motion tracking system: LeVan et al., 2013), which do not require R-peak detections for the BCG artefact corrections, might be alternatives to the AAS approach. These hardware approaches would especially be suited in an ultra-high field MRI scanner such as 7T MRI, as the ECG distortion magnifies and becomes more problematic as compared to 3T MRI (Brookes et al., 2009; Debener et al., 2008; Jorge et al., 2019, 2015a, 2015b; Mullinger et al., 2008a; Wirsich et al., 2021). Moving away from the techniques that requires R-peak detection from the ECG signals, as suggested by our present work, could provide a more reliable and consistent way to process and analyse EEG-fMRI data.

### Limitations

In this study, we demonstrated that this beamforming approach did not only attenuate the BCG artefacts, but also preserved the identified spindle activity even when compared to the conventional AAS BCG corrections. However, our approach also has some limitations. For the construction of the beamforming spatial filters, realistic head volume conductor modelling is required for accurately computing the EEG and MEG lead-fields (Brookes et al., 2008a; Neugebauer et al., 2017). Although accurate head models based on realistic head geometry obtained from individual anatomical MR images and knowledge of the accurate location of the EEG electrodes on the head can provide a more accurate EEG forward solution, access to such information might not always be available. In this study, we did consider a realistic head model, consisting of a four-layer (scalp, skull, CSF, and brain) BEM head model from each individual anatomical images, with digitized EEG electrode locations using the EGI GPS to facilitate individualized accurate co-registration of electrode positions with each subject’s anatomical image. Without these kinds of information, the beamforming approach might be suboptimal.

Another limitation is that the use of high-density EEG electrodes (at least 64 EEG channels) might be required to better suppress the MRI-related artefacts. Brookes et al. (2009) demonstrated that, when the data was acquired at 7T MRI, purely increasing the number of EEG channels from 32 to 64 allowed them to improve the SNRs by a factor of around 1.6 by attenuating the MRI-related artefacts. Another major limitation of beamforming technique is that this approach cannot properly reconstruct two spatially separate but temporally correlated sources (Brookes et al., 2007; Huang et al., 2004; Quraan and Cheyne, 2010; van Veen et al., 1997). For example, beamforming could cancel two temporally correlated sources when they are spatially far from each other (i.e., auditory steady-state response) (Brookes et al., 2007) or merge them when they are spatially close to each other (Huang et al., 2004). It is worth mentioning that a modified source model instead of a standard single source model allows for beamforming reconstruction of the spatially separate but temporally correlated sources (Brookes et al., 2007; Diwakar et al., 2011; Kuznetsova et al., 2021).

It should also be considered to optimize beamforming weights for spatial filtering for the whole sleep recording. In this study, we conducted our analysis based on the spindle events detected by expert sleep scorers, and used the baseline periods before each spindle onset to optimize the beamforming spatial filter. Since the beamforming approach is beneficial to distinguish sleep related EEG activity and the MRI-related artefacts, it would be crucial to consider this approach even for sleep staging. However, lack of clear baselines and different vigilant stages (i.e., awake, N1, N2, N3, REM) in the sleep recording might result in a suboptimal beamforming spatial filter to supress the artefacts for the entire sleep recording. Although the beamforming approach has been used for the resting-state data in MEG (Hillebrand et al., 2016, 2012), certain time-windows (around 290s or around 265s in respective papers) of the same vigilant state (i.e., awake resting-state) has been used to compute the data covariance matrix for the beamforming weights. Furthermore, EEG signals during the EEG-fMRI are also noisier when compared to MEG recordings. It might be interesting to investigate the behaviours of the beamforming approach to reconstruct time-courses of the whole sleep recording during EEG-fMRI. An optimal beamforming spatial filtering that suppresses the MRI-related noise across different vigilant states and preserves the sleep related EEG signals would allow more confident sleep staging of EEG-fMRI data on par with the sleep staging outside the MRI scanner.

### Implications

Since this study aimed at investigating the benefits of beamforming approach in the EEG-fMRI analysis acquired during sleep, we have limited our analysis to typical spindle features (11-16Hz) instead of specifically separating fast (12-15 Hz) and slow (9-12 Hz) spindles. Confirmed by our findings, our approach could be used to better differentiate between the fast and slow spindles in EEG-fMRI. Previous literature demonstrated the distinct characteristics of the fast and slow spindles in terms of occurrence and topography (Cox et al., 2017; Mölle et al., 2011). Especially the fast spindles are expected to play a key role in sleep-dependent memory processing (Fogel and Smith, 2011; Jegou et al., 2019; Mölle et al., 2011). In addition to the role of memory consolidations, spindles have also been proposed to participate in selectively processing and suppressing external stimuli, such as auditory tones, during sleep (Dang-Vu et al., 2011, 2010a; Schabus et al., 2012). This mechanism is known as a protective mechanism both maintaining sleep and ensuring quality of sleep. Our proposed approach might help better distinguish and characterise different types of spindles for those different functions. Although we have not investigated slow waves in this study, we believe that this approach could be applied to further advance our understanding of slow wave activity. Lastly, more people currently suffer from short or disturbed sleep, and sleep disorders (i.e., insomnia, hypersomnia) have been steadily increasing in our society. In order to better understand and characterize such sleep disorders, EEG-fMRI is a promising method, and our proposed approach would provide a new insight by ensuring to differentiate sleep-related brain activities and the MRI-related artefacts.

## Conclusions

We demonstrated that our data-driven beamforming approach did not only attenuate the BCG artefacts without the need for ECG R-peak detection, but also recovered sleep spindle activity normally occurring in NREM sleep, which the BCG artefacts typically obscure. Our findings support and extend the benefit of applying beamforming source imaging technique to the EEG-fMRI acquired during sleep in order to extract meaningful sleep related brain signals and suppress the MRI-related artefacts. We demonstrated that this approach would be beneficial especially for long EEG-fMRI data acquisitions (i.e., sleep, resting-state) when it is more time consuming to semi-automatically detect cardiac events. Furthermore, using this approach allowed us to accurately estimate single-event power change of each spindle in the source space when compared to the channel level, and to improve the specificity of fMRI GLM analysis by better localizing the recruited brain regions during spindles as indicated by reduced number of significant voxels both in total and outside of the gray matter regions. Future research could use this approach to better characterize the role of spindles (i.e., fast vs. slow spindles) for memory consolidations and protective mechanisms during sleep. Lastly, this approach might be useful to better investigate neurovascular couplings of those suffering from sleep disorders (i.e., insomnia, hypersomnia) by ensuring to differentiate sleep-related brain activities and the MRI-related artefacts.

## The authorship contribution statement

**Makoto Uji**: Conceptualization, Methodology, Formal data analysis, Investigation, Data curation, Writing - original draft, Visualization, Project administration. **Aude Jegou**: Data acquisition, Writing - review & editing. **Nathan Cross**: Data acquisition, Sleep scoring, Spindle analysis, Writing - review & editing. **Florence B. Pomares**: Data acquisition, Writing - review & editing. **Aurore A. Perrault**: Data acquisition, Sleep scoring, Spindle analysis, Writing - review & editing. **Alex Nguyen**: Data acquisition, Writing - review & editing. **Umit Aydin**: Data acquisition, Writing - review & editing. **Kangjoo Lee**: Data acquisition, Writing - review & editing. **Chifaou Abdallah**: Data acquisition, Sleep scoring, Spindle analysis, Writing - review & editing. **Birgit Frauscher:** Methodology, Sleep scoring, Spindle analysis, Writing - review & editing. **Jean-Marc Lina**: Methodology, Validation, Writing - review & editing. **Thien Thanh Dang-Vu**: Conceptualization, Resources, Writing - review & editing, Supervision, Project administration, Funding acquisition. **Christophe Grova**: Conceptualization, Resources, Writing - review & editing, Supervision, Project administration, Funding acquisition.

## Acknowledgements

This research was supported by the Natural Sciences and Engineering Research Council of Canada (TDV) and the Canada Foundation for Innovation (TDV). The MRI compatible high-density EEG device (Magstim-EGI) and data acquisition were made possible through an internal grant from PERFORM centre and the Faculty of Arts and Science of Concordia University (CG). TDV is also supported by the Canadian Institutes of Health Research (MOP 142191, PJT 153115, PJT 156125 and PJT 166167), the Fonds de Recherche du Québec – Santé and Concordia University. CG is supported by Natural Sciences and Engineering Research Council of Canada Discovery grants as well as the Canadian Institutes of Health Research (PJT-159948 and MOP-133619) and the Fonds de Recherche du Québec, Nature et Technology (research team grant). MU is supported by a Horizon Postdoctoral Fellowship from Concordia University. BF is supported by a salary award (“Chercheur-boursier clinicien Senior”) of the Fonds de Recherche du Québec – Santé 2021-2025, a NSERC discovery grant and accelerator supplement 2020-2025, CIHR (PJT-175056) 2021-2026, and the Hewitt Foundation 2019-2026.

## Supplementary

**Figure S1.**
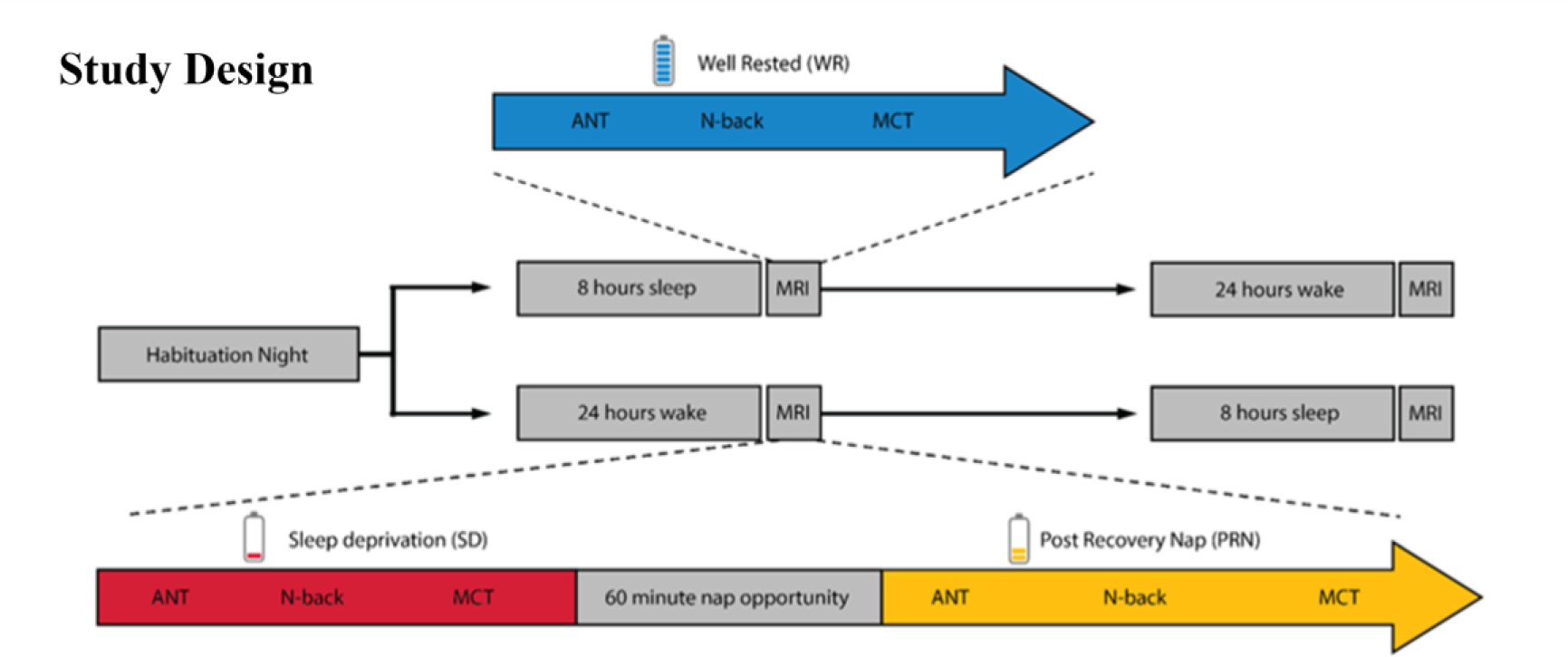
Experimental study design (Cross et al, 2021; 2021). Participants made 3 visits to the lab: a habituation night, followed by a counterweighted design of either another full-night opportunity to sleep (blue), or a night of total sleep deprivation. In the morning following each night, participants completed a resting-state and 3 cognitive tasks (Attentional Network Task: ANT, Mackworth clock task: MCT, N-back task) inside the MRI scanner. In the sleep deprived state (red), participants also had a recovery nap opportunity (yellow) and then repeated the resting-state and tasks inside the MRI scanner.

**Figure S2.**
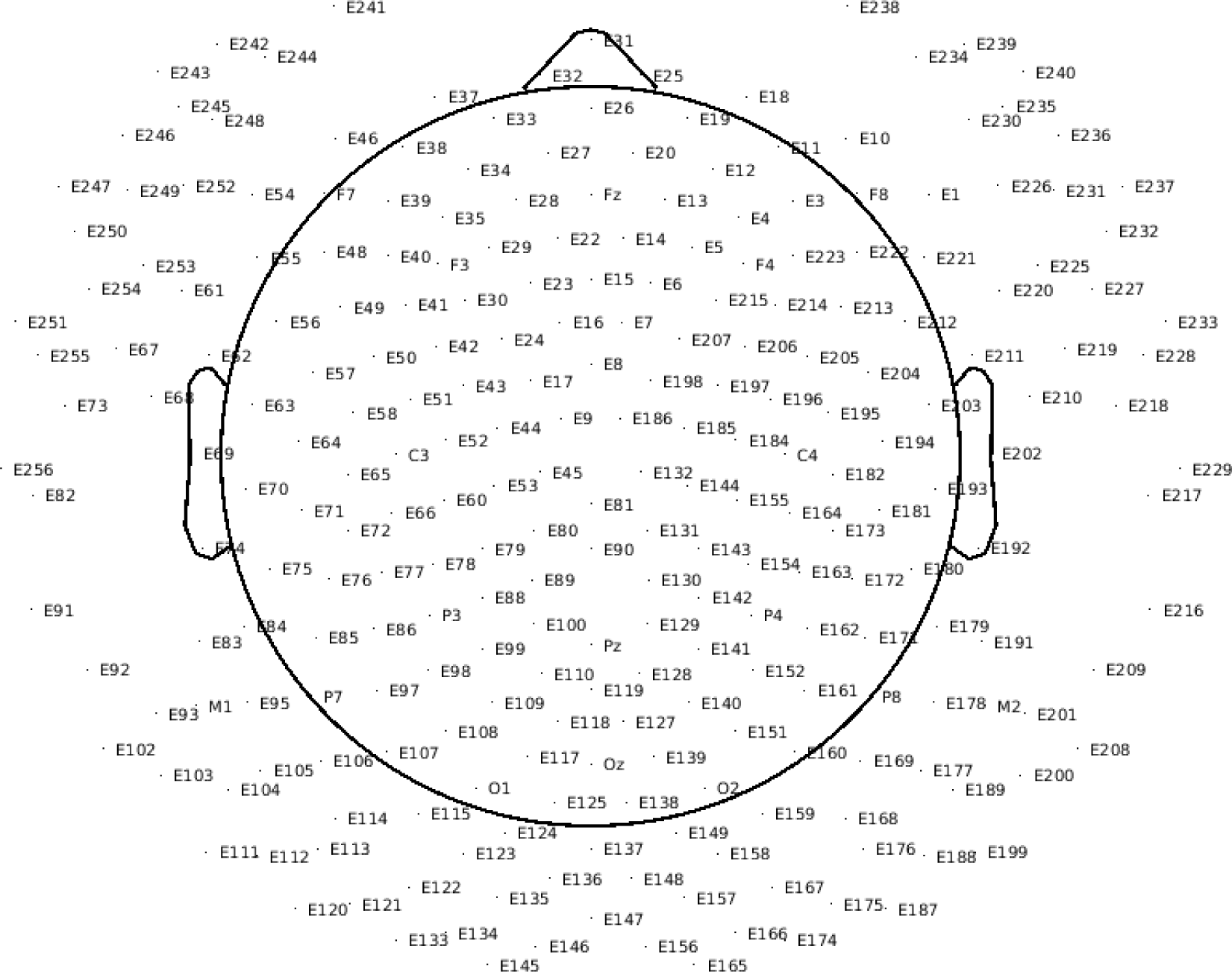
256 channel EGI EEG 2D layout. Two lowest rows of EEG electrodes around the neck and face (69 electrodes: E73, E82, E91, E92, E93, E102, E103, E104, E111, E112, E113, E120, E121, E122, E133, E134, E135, E145, E146, E147, E156, E157, E165, E166, E167, E174, E175, E176, E187, E188, E189, E199, E200, E201, E208, E209, E216, E217, E218, E226, E227, E228, E229, E230, E231, E232, E233, E234, E235, E236, E237, E238, E239, E240, E241, E242, E243, E244, E245, E246, E247, E248, E249, E250, E251, E252, E254, E255, E256) were removed keeping 187 electrodes for data analysis.

**Figure S3.**
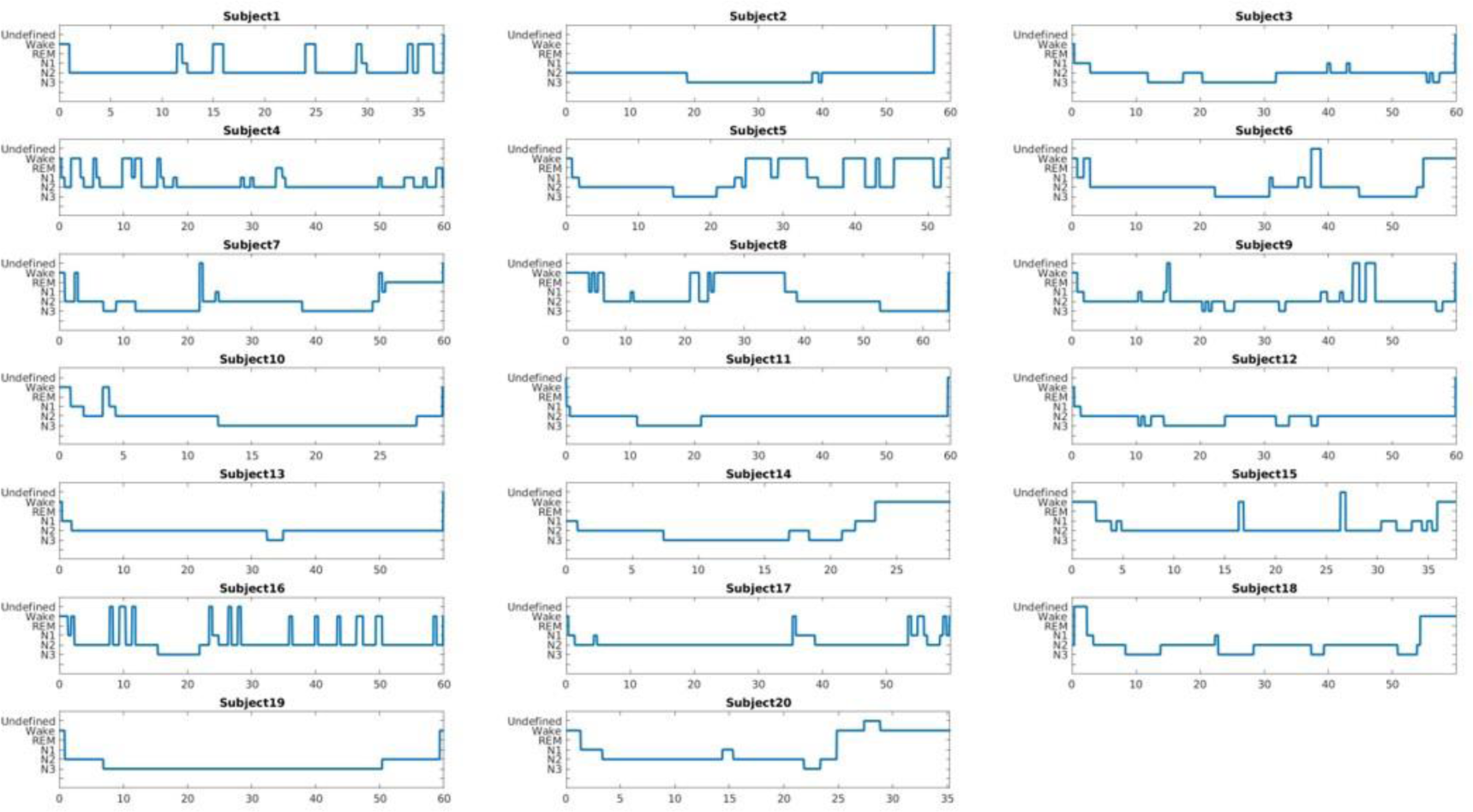
20 subjects’ sleep hypnograms during the whole nap EEG-fMRI session. X axis displays time in minutes.

**Figure S4.**
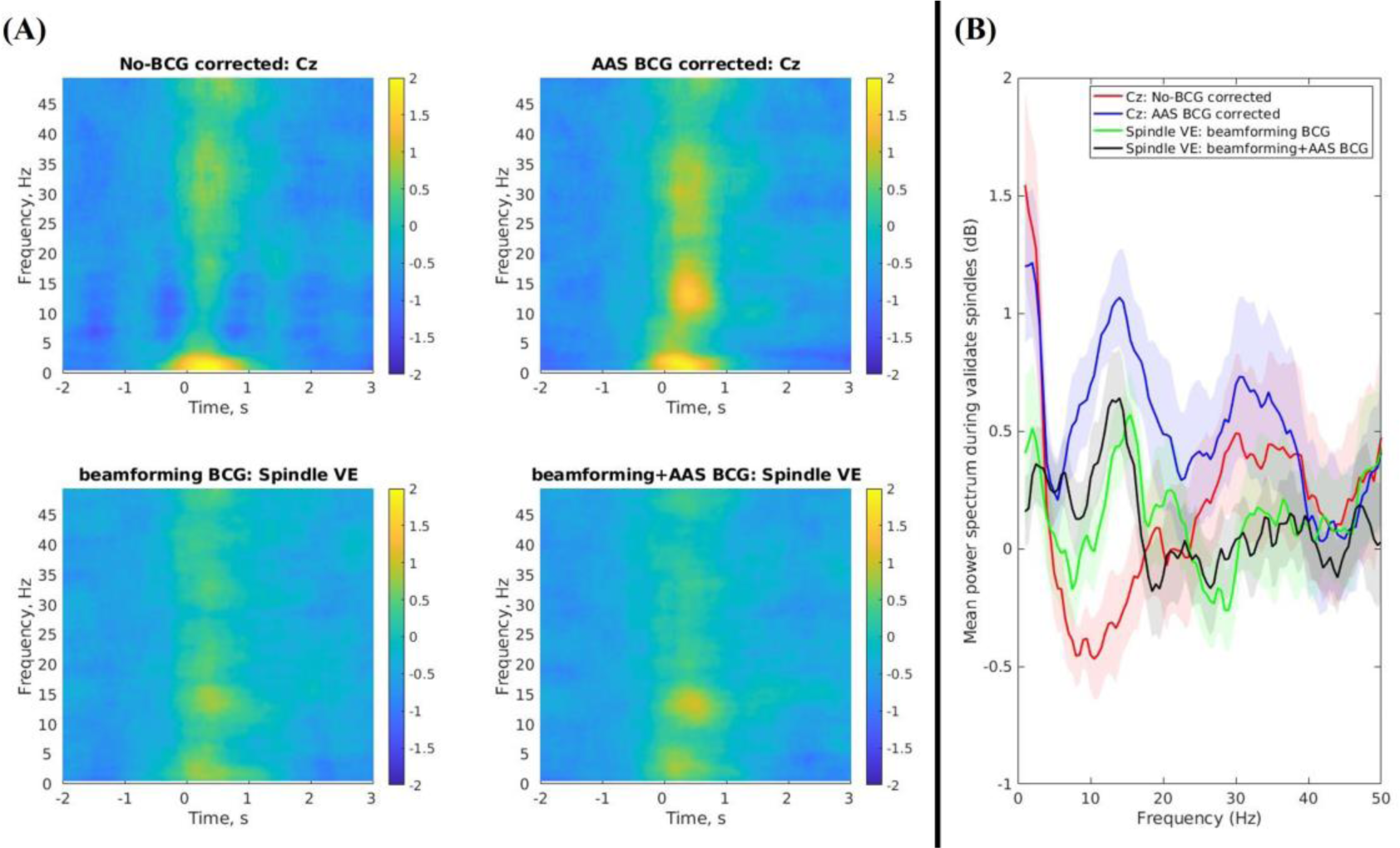
**(A)** Group mean (N=20) time-frequency representations (TFRs) of spindle activity during the validated NREM periods for the non-BCG corrected (Cz: top left), AAS BCG corrected (Cz: top right), beamforming BCG corrected (spindle VE: bottom left), and beamforming+AAS BCG corrected data (spindle VE: bottom right). These TFRs demonstrated −2s to 3s relative to the spindle onsets (t=0). **(B)** Group mean (N=20) power spectrums of the validated spindle durations during the validated NREM periods (M±SD = 0.84±0.06s).

**Figure S5.**
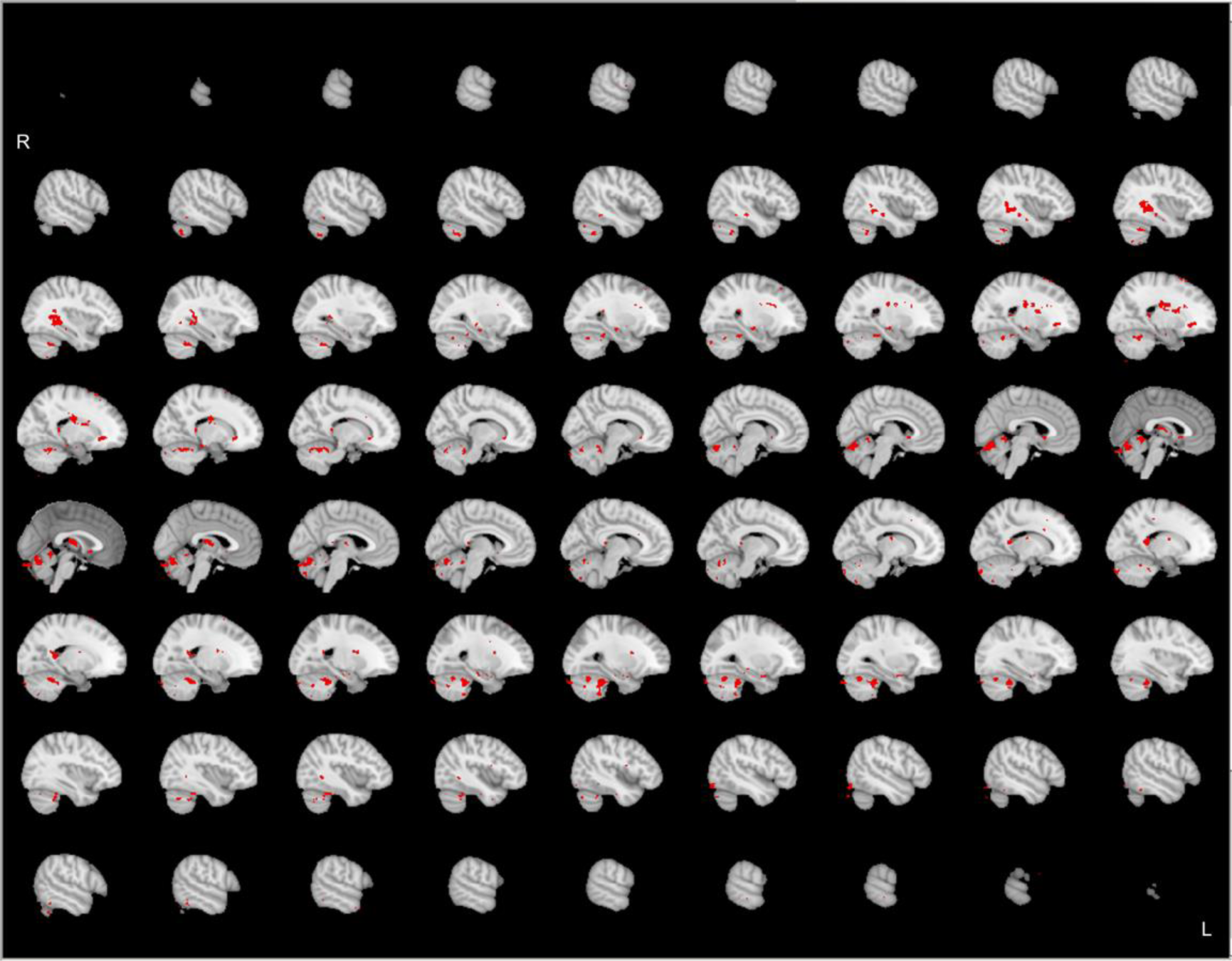
Group-level (N=20) significant main effects for GLM1 (spindle onset & duration) during the validated NREM periods, revealed by FSL randomize non-parametric permutation testing with 5,000 permutations (Eklund et al., 2016; Winkler et al., 2014) with the voxel-wise inference (p < 0.05 corrected). This statistical map displays at uncorrected p < 0.001.

**Figure S6.**
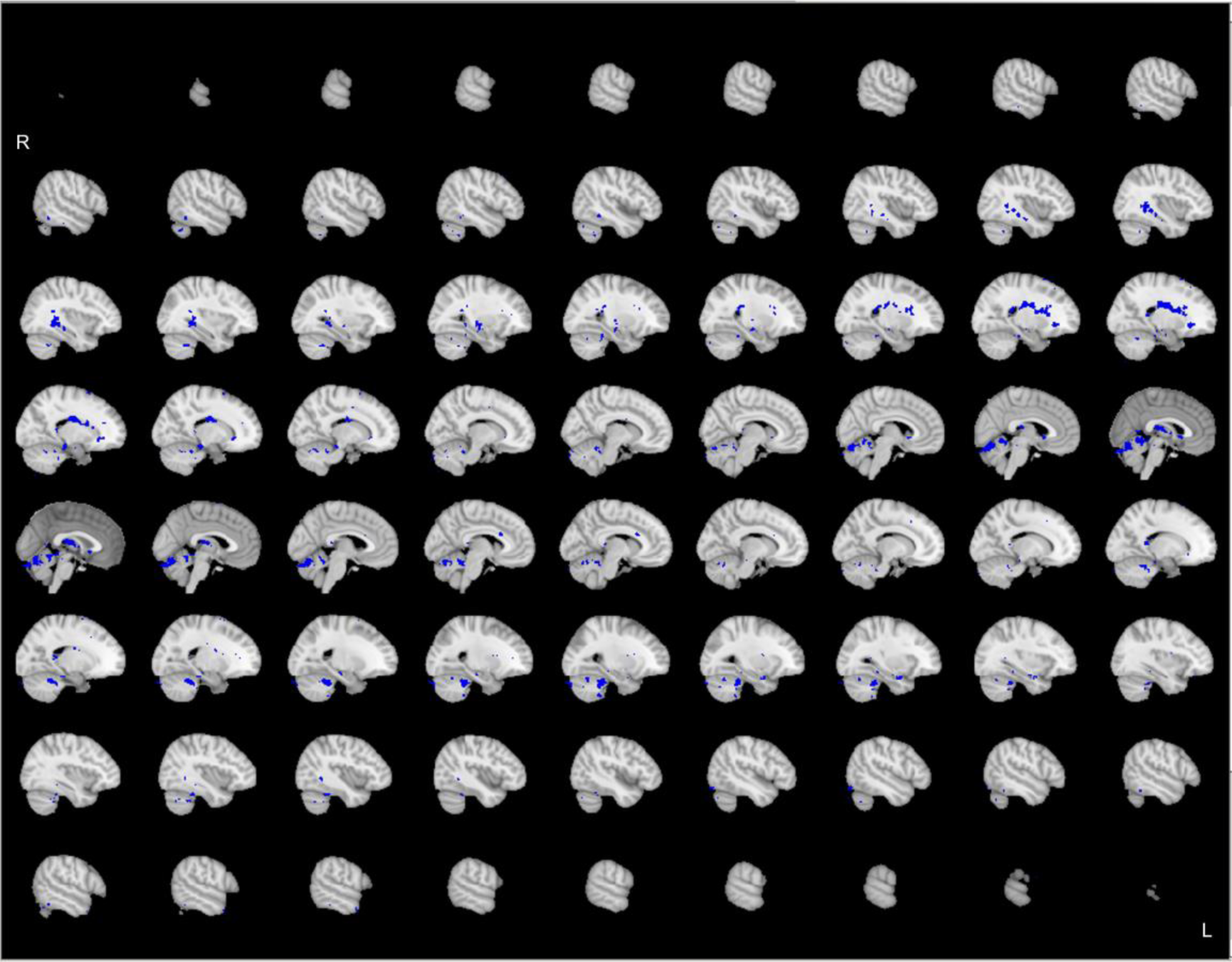
Group-level (N=20) significant main effects for GLM2 (spindle onset, duration, & single-spindle power changes from the Cz electrode of the AAS BCG corrected data) during the validated NREM periods, revealed by FSL randomize non-parametric permutation testing with 5,000 permutations (Eklund et al., 2016; Winkler et al., 2014) with the voxel-wise inference (p < 0.05 corrected). This statistical map displays at uncorrected p < 0.001.

**Figure S7.**
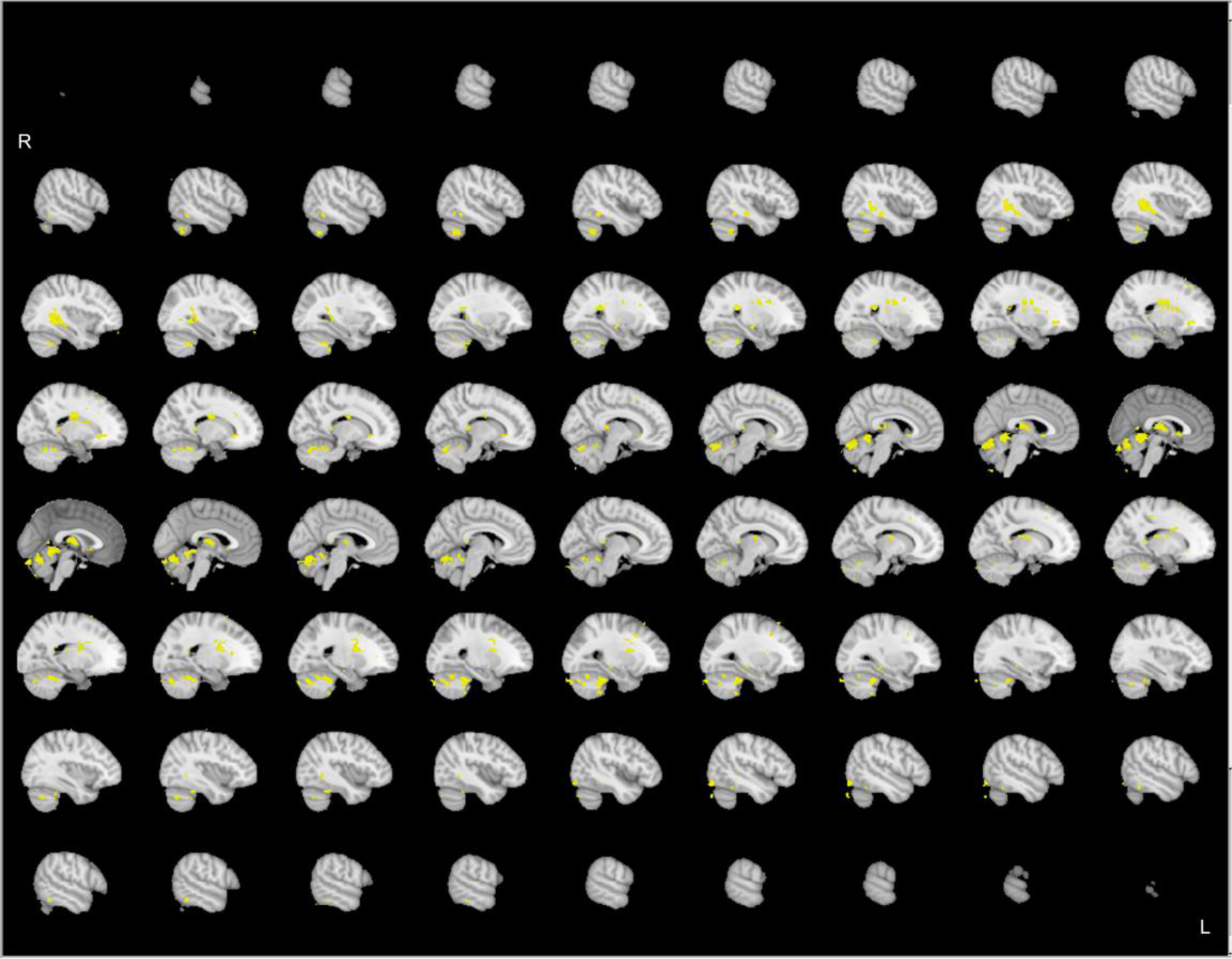
Group-level (N=20) significant main effects for GLM3 (spindle onset, duration, & single-spindle power changes from the spindle VE of the beamforming+AAS BCG corrected data) during the validated NREM periods, revealed by FSL randomize non-parametric permutation testing with 5,000 permutations (Eklund et al., 2016; Winkler et al., 2014) with the voxel-wise inference (p < 0.05 corrected). This statistical map displays at uncorrected p < 0.001.

**Figure S8.**
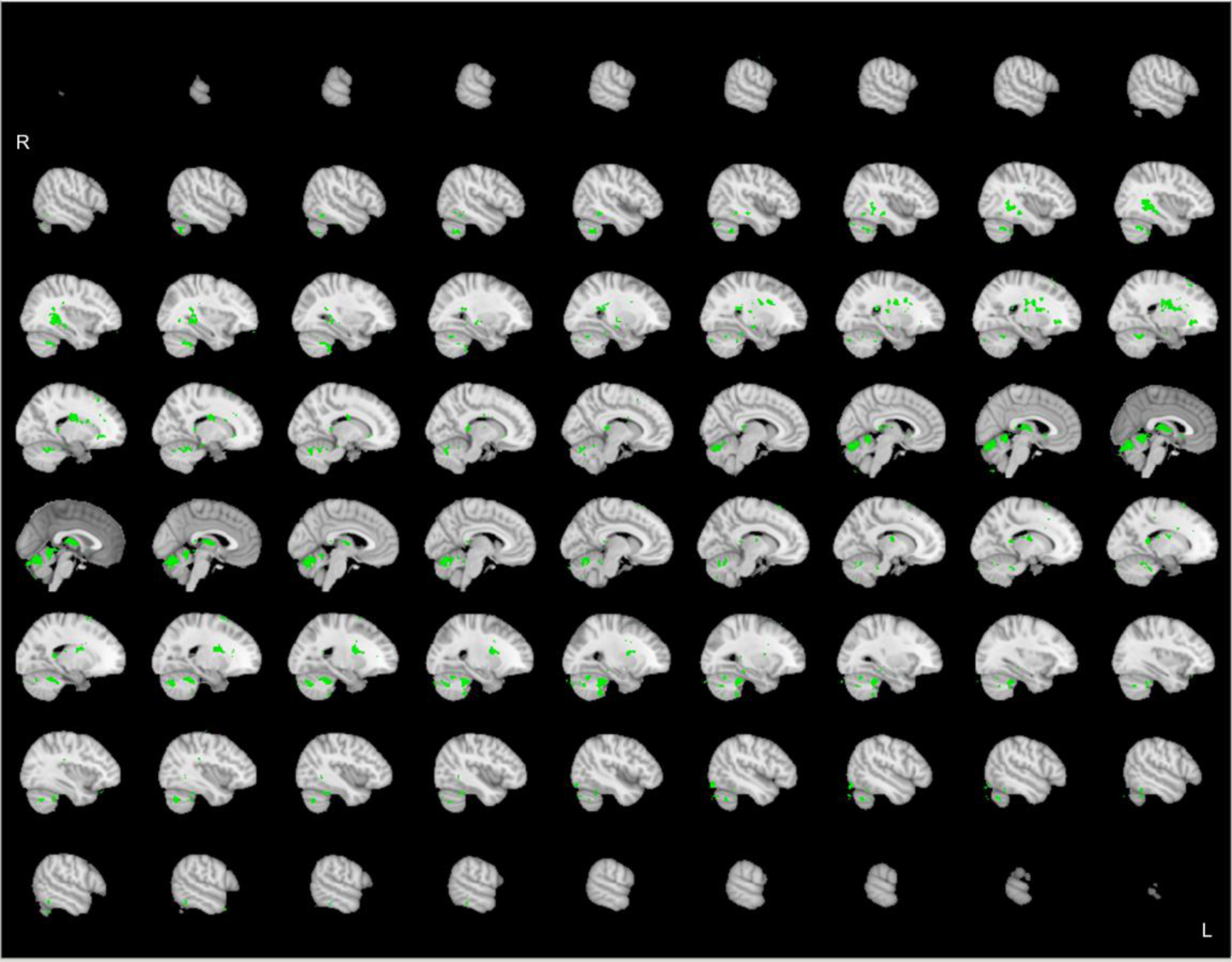
Group-level (N=20) significant main effects for GLM4 (spindle onset, duration, & single-spindle power changes from the spindle VE of the beamforming BCG corrected data) during the validated NREM periods, revealed by FSL randomize non-parametric permutation testing with 5,000 permutations (Eklund et al., 2016; Winkler et al., 2014) with the voxel-wise inference (p < 0.05 corrected). This statistical map displays at uncorrected p < 0.001.

**Figure S9.**
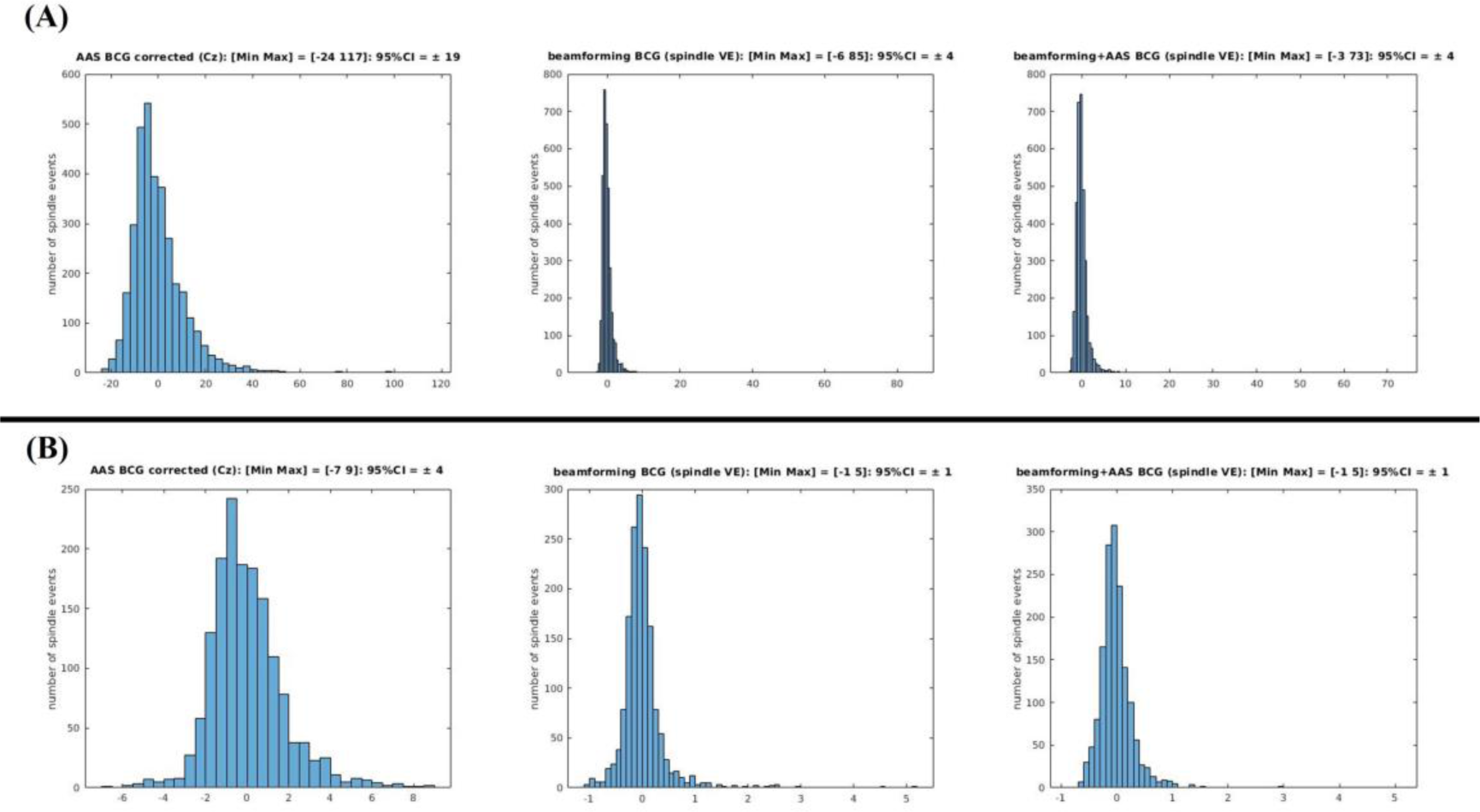
Spindle event power amplitude variability across 20 subjects during **(A) the whole nap** and **(B) the validated NREM periods.** Left figures show the spindle amplitude variability from average artefact subtraction (AAS) method ballistocardiogram (BCG) corrected data (Cz: left), middle figures show those from beamforming BCG corrected data (spindle VE: middle), and right figures show beamforming+AAS BCG corrected (spindle VE: right). 95% confident intervals of the channel level spindle power amplitude (AAS BCG corrected) were ±19 (A.U.) during the whole nap and ±4 (A.U.) during the validated NREM periods, whereas those in both source levels (beamforming BCG corrected; beamforming+AAS BCG corrected) were ±4 during the whole nap and ±1 (A.U.) during the validated NREM periods for both source level corrections. These amplitude variabilities were extracted by demeaning each subject’s Hilbert transformed spindle frequency (11-16Hz) signals during each spindle duration. These larger amplitude changes in the channel-level might be due to the residual BCG artefacts causing less accurate spindle power change estimates for the GLM regressors when compared to the source level estimates (GLM3 & GLM4) or just a boxcar ([1 0], GLM1) regressor.

