## Supplemental for "EEG source imaging technique to investigate sleep oscillations for simultaneous EEG-fMRI"

**Supplementary**

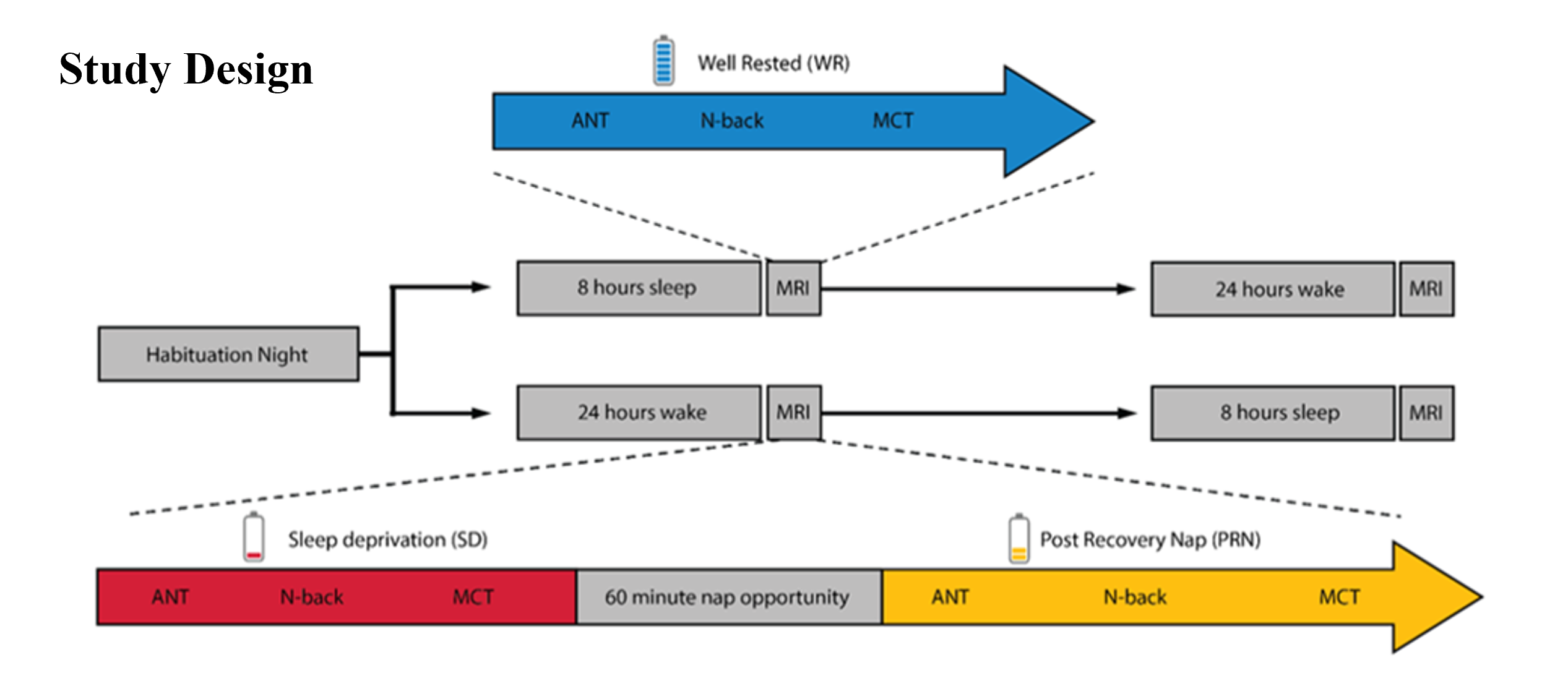

**Figure S1.** Experimental study design (Cross et al, 2021; 2021). Participants made 3 visits to the lab: a habituation night, followed by a counterweighted design of either another full-night opportunity to sleep (blue), or a night of total sleep deprivation. In the morning following each night, participants completed a resting-state and 3 cognitive tasks (Attentional Network Task: ANT, Mackworth clock task: MCT, N-back task) inside the MRI scanner. In the sleep deprived state (red), participants also had a recovery nap opportunity (yellow) and then repeated the resting-state and tasks inside the MRI scanner.

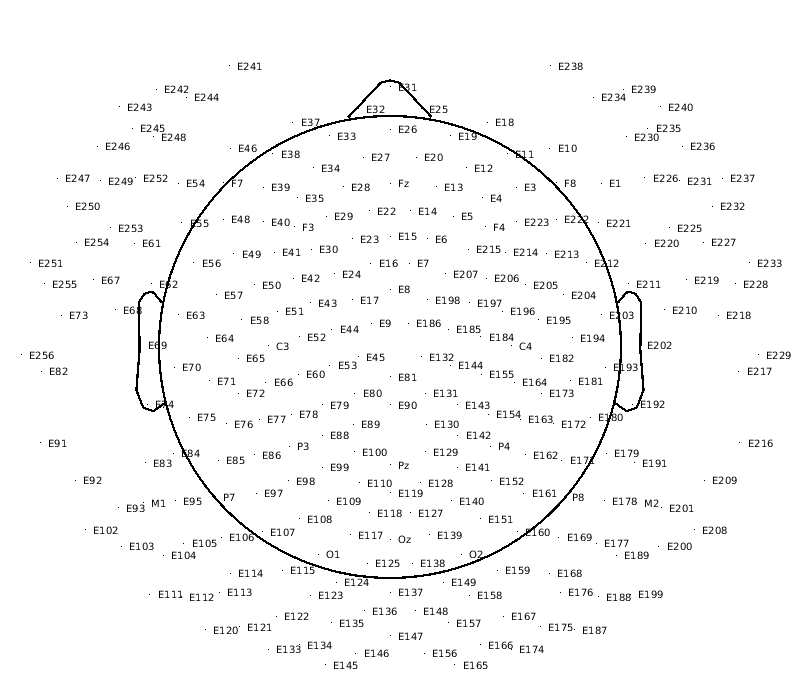

**Figure S2.** 256 channel EGI EEG 2D layout. Two lowest rows of EEG electrodes around the neck and face (69 electrodes: E73, E82, E91, E92, E93, E102, E103, E104, E111, E112, E113, E120, E121, E122, E133, E134, E135, E145, E146, E147, E156, E157, E165, E166, E167, E174, E175, E176, E187, E188, E189, E199, E200, E201, E208, E209, E216, E217, E218, E226, E227, E228, E229, E230, E231, E232, E233, E234, E235, E236, E237, E238, E239, E240, E241, E242, E243, E244, E245, E246, E247, E248, E249, E250, E251, E252, E254, E255, E256) were removed keeping 187 electrodes for data analysis.

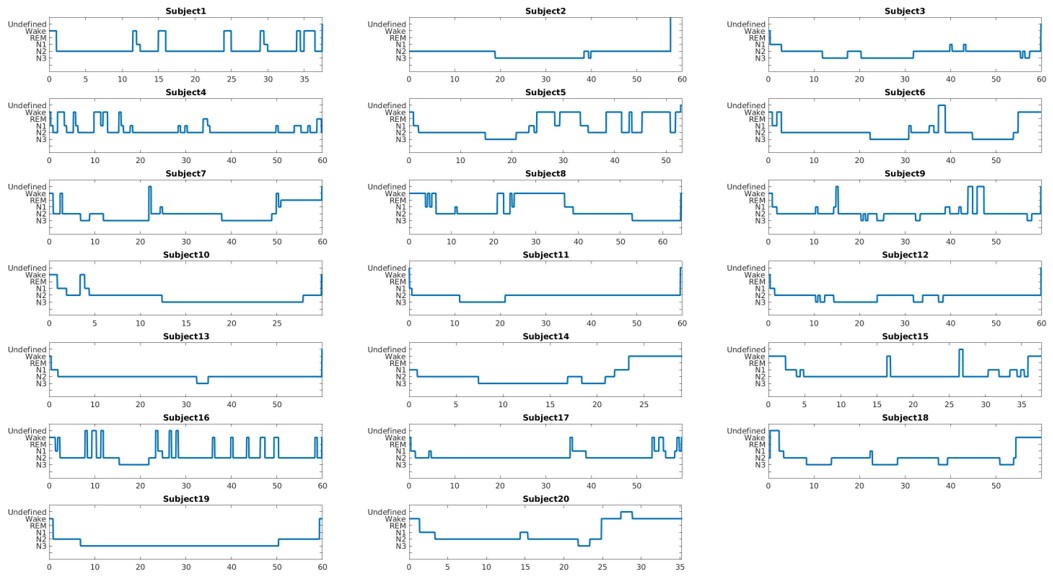

**Figure S3.** 20 subjects’ sleep hypnograms during the whole nap EEG-fMRI session. X axis displays time in minutes.

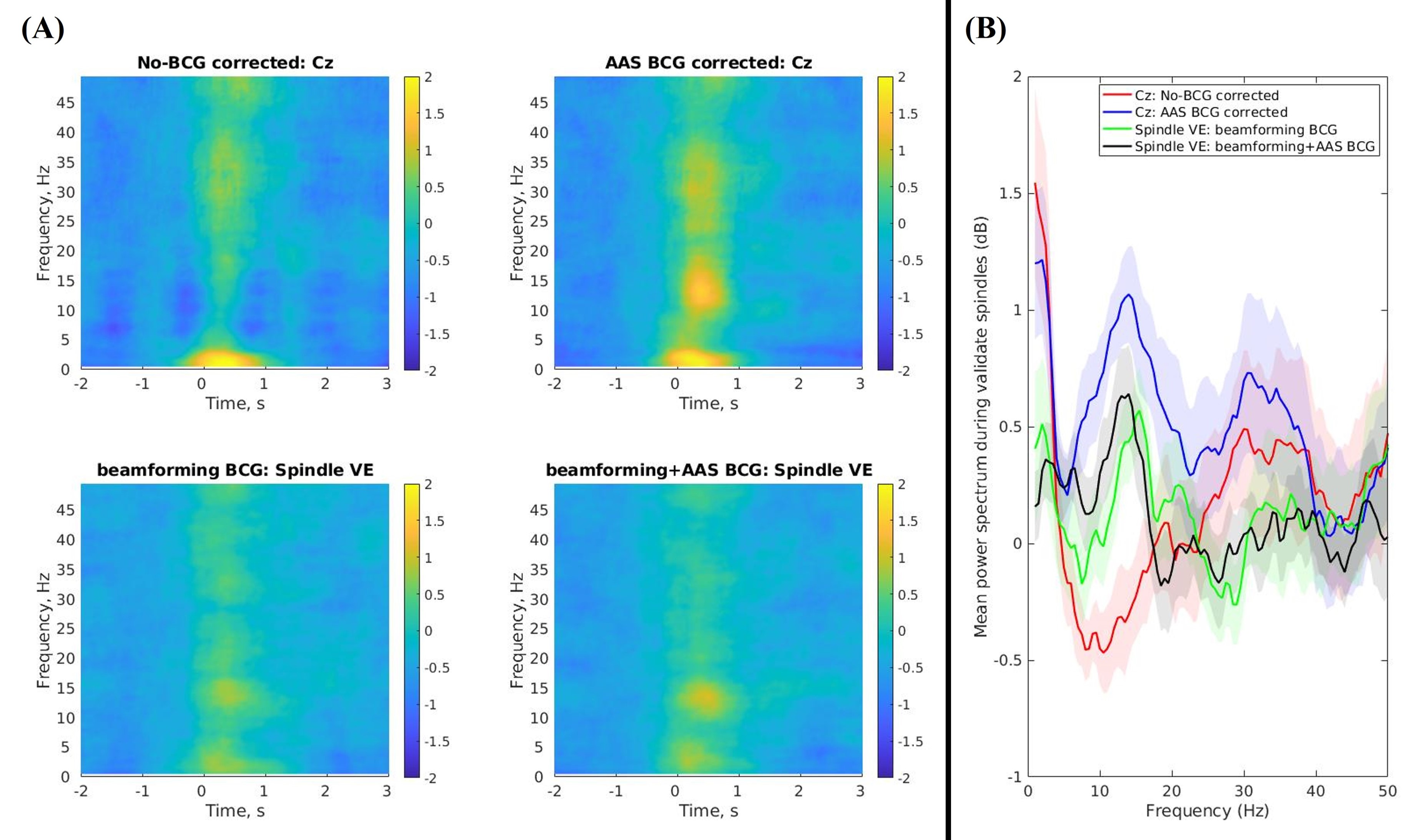

**Figure S4. (A)** Group mean (N=20) time-frequency representations (TFRs) of spindle activity during the validated NREM periods for the non-BCG corrected (Cz: top left), AAS BCG corrected (Cz: top right), beamforming BCG corrected (spindle VE: bottom left), and beamforming+AAS BCG corrected data (spindle VE: bottom right). These TFRs demonstrated -2s to 3s relative to the spindle onsets (t=0). **(B)** Group mean (N=20) power spectrums of the validated spindle durations during the validated NREM periods (M±SD = 0.84±0.06s).

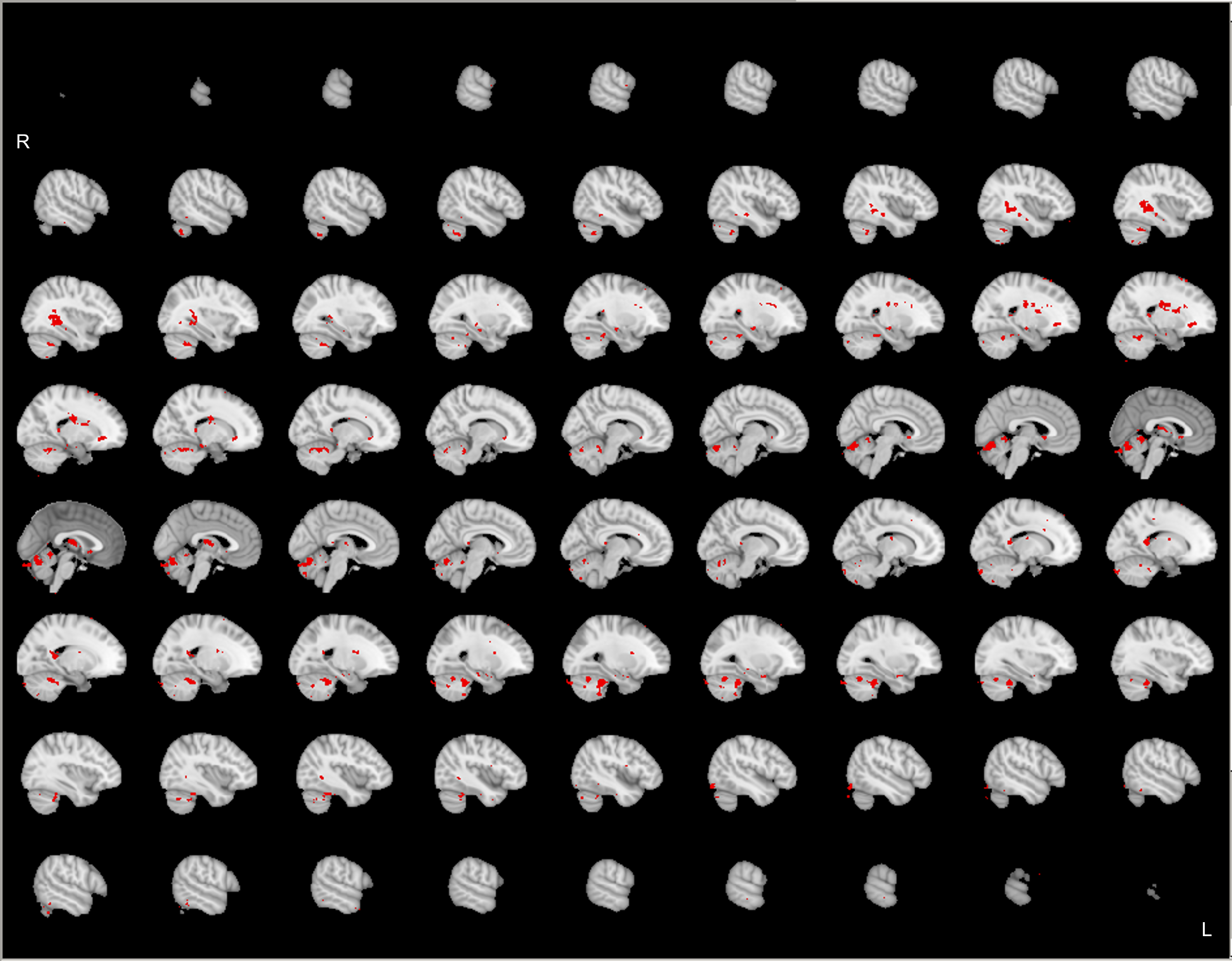

**Figure S5.** Group-level (N=20) significant main effects for GLM1 (spindle onset & duration) during the validated NREM periods, revealed by FSL randomize non-parametric permutation testing with 5,000 permutations (Eklund et al., 2016; Winkler et al., 2014) with the voxel-wise inference (p < 0.05 corrected). This statistical map displays at uncorrected p < 0.001.

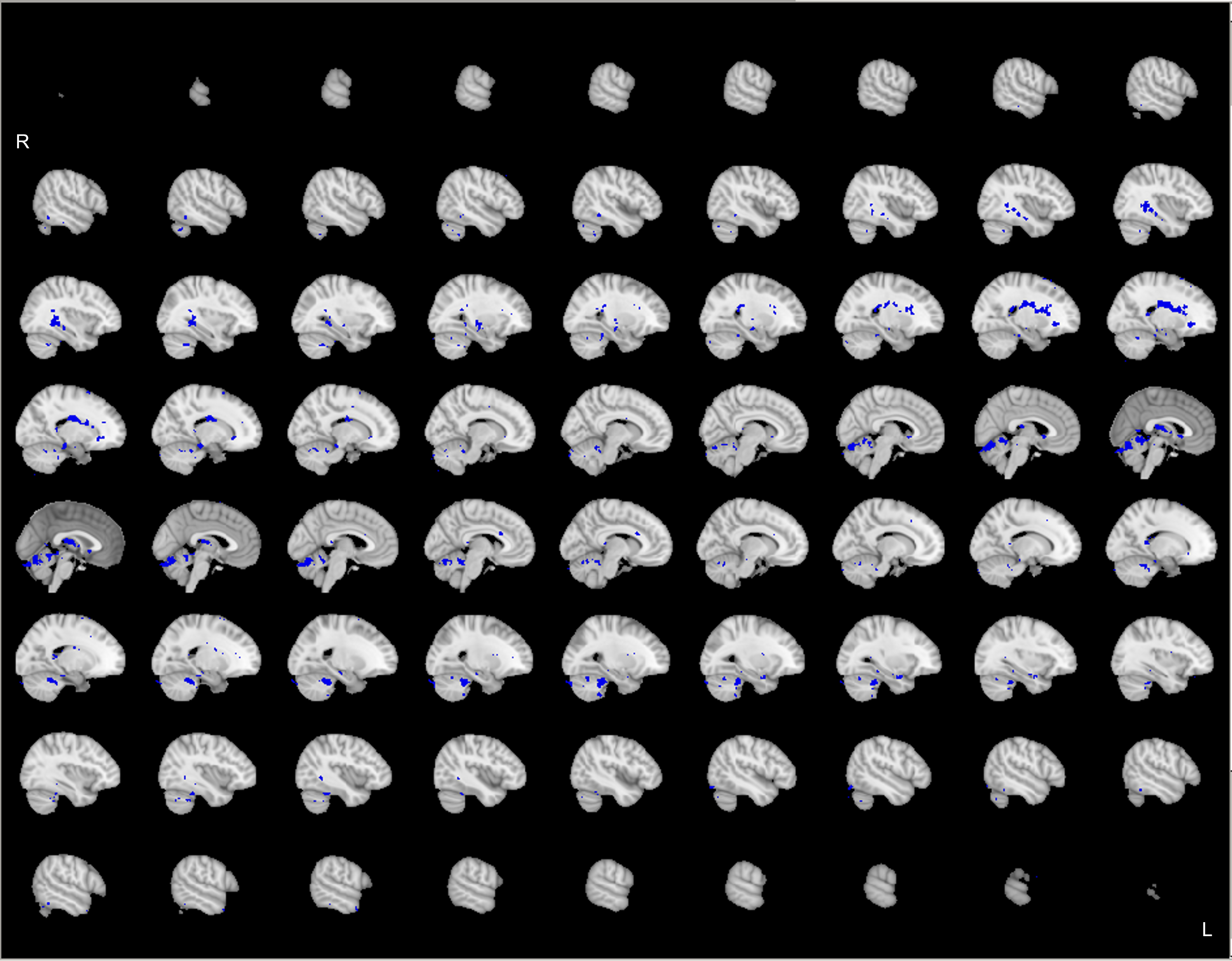

**Figure S6.** Group-level (N=20) significant main effects for GLM2 (spindle onset, duration, & single-spindle power changes from the Cz electrode of the AAS BCG corrected data) during the validated NREM periods, revealed by FSL randomize non-parametric permutation testing with 5,000 permutations (Eklund et al., 2016; Winkler et al., 2014) with the voxel-wise inference (p < 0.05 corrected). This statistical map displays at uncorrected p < 0.001.

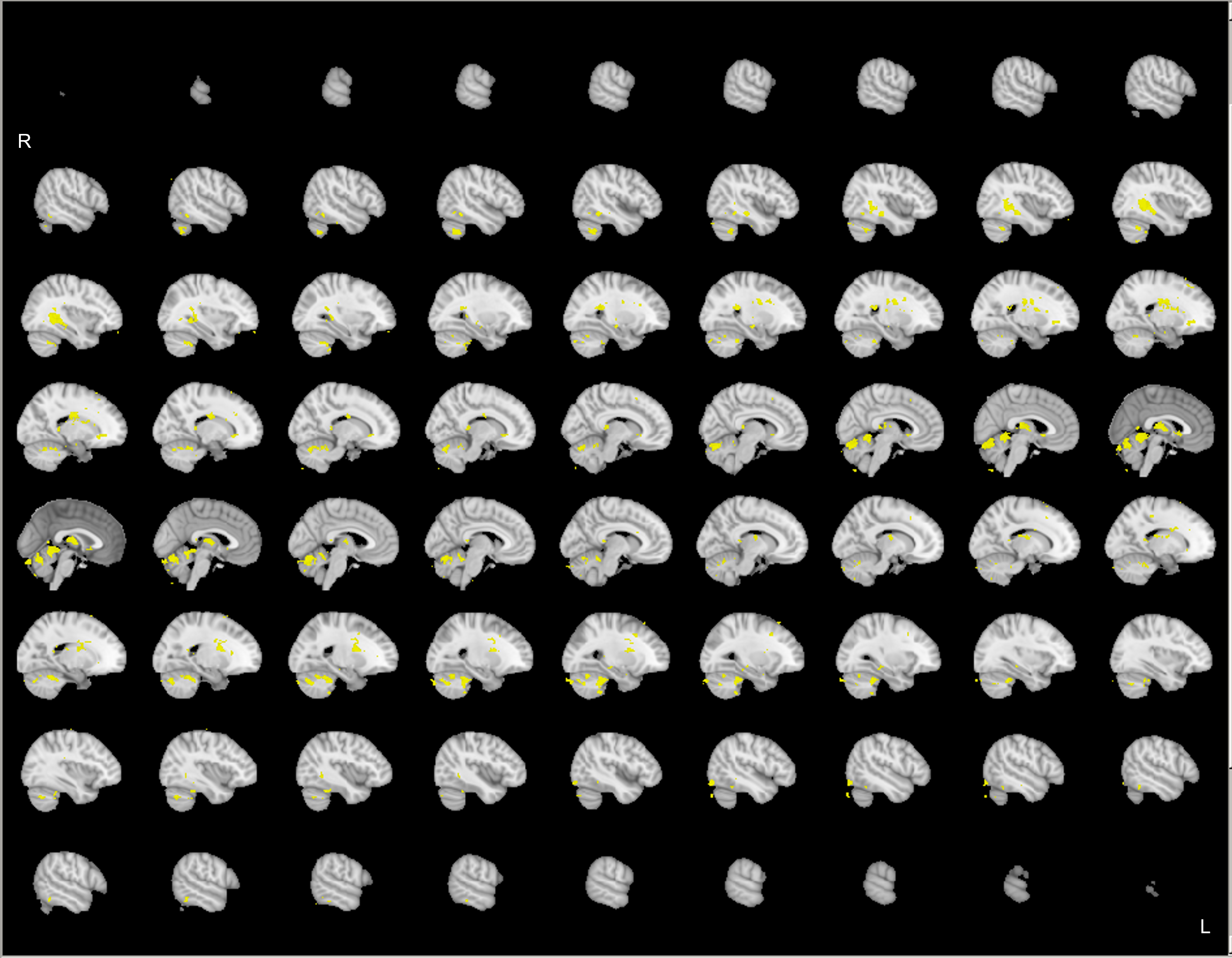

**Figure S7.** Group-level (N=20) significant main effects for GLM3 (spindle onset, duration, & single-spindle power changes from the spindle VE of the beamforming+AAS BCG corrected data) during the validated NREM periods, revealed by FSL randomize non-parametric permutation testing with 5,000 permutations (Eklund et al., 2016; Winkler et al., 2014) with the voxel-wise inference (p < 0.05 corrected). This statistical map displays at uncorrected p < 0.001.

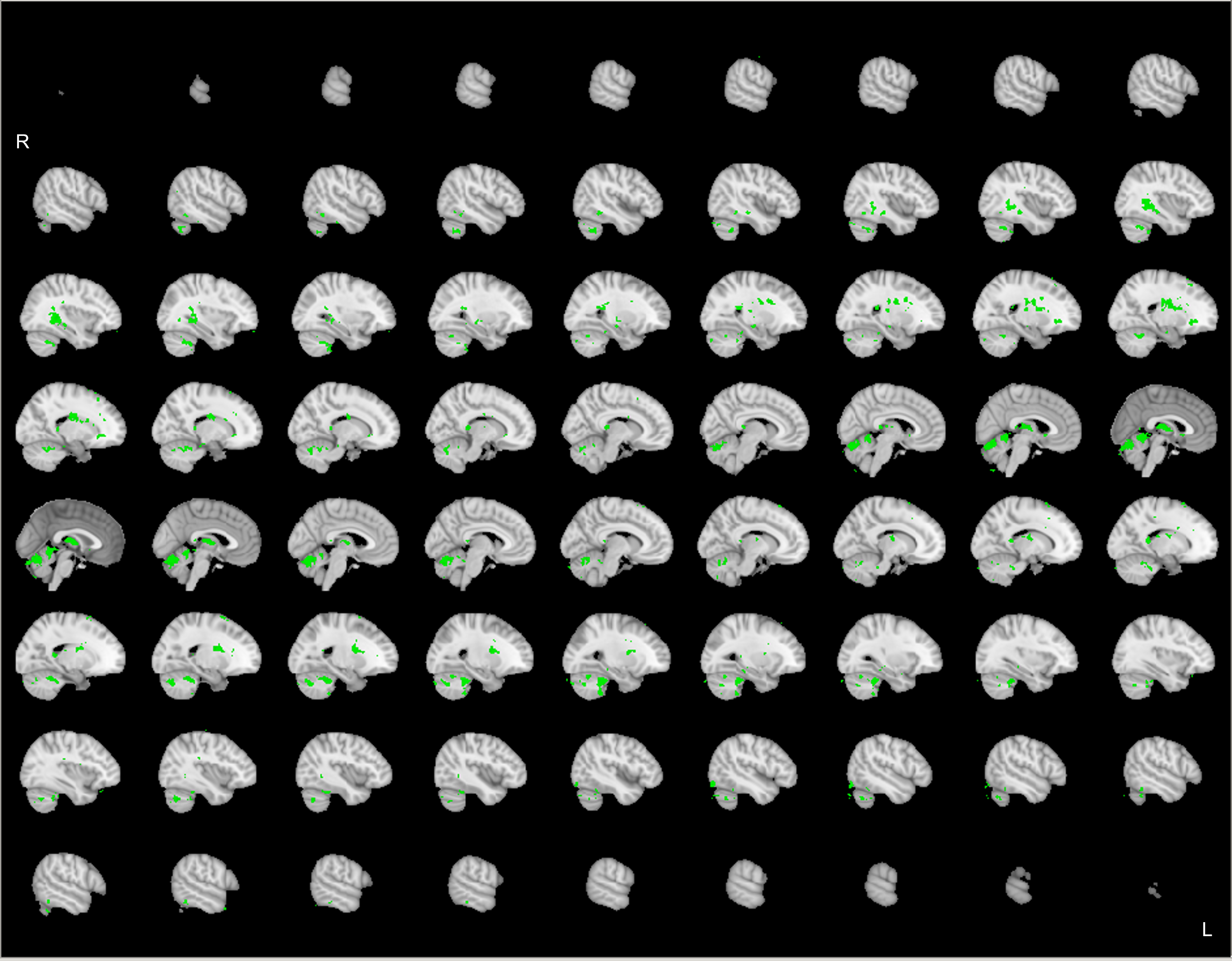

**Figure S8.** Group-level (N=20) significant main effects for GLM4 (spindle onset, duration, & single-spindle power changes from the spindle VE of the beamforming BCG corrected data) during the validated NREM periods, revealed by FSL randomize non-parametric permutation testing with 5,000 permutations (Eklund et al., 2016; Winkler et al., 2014) with the voxel-wise inference (p < 0.05 corrected). This statistical map displays at uncorrected p < 0.001.

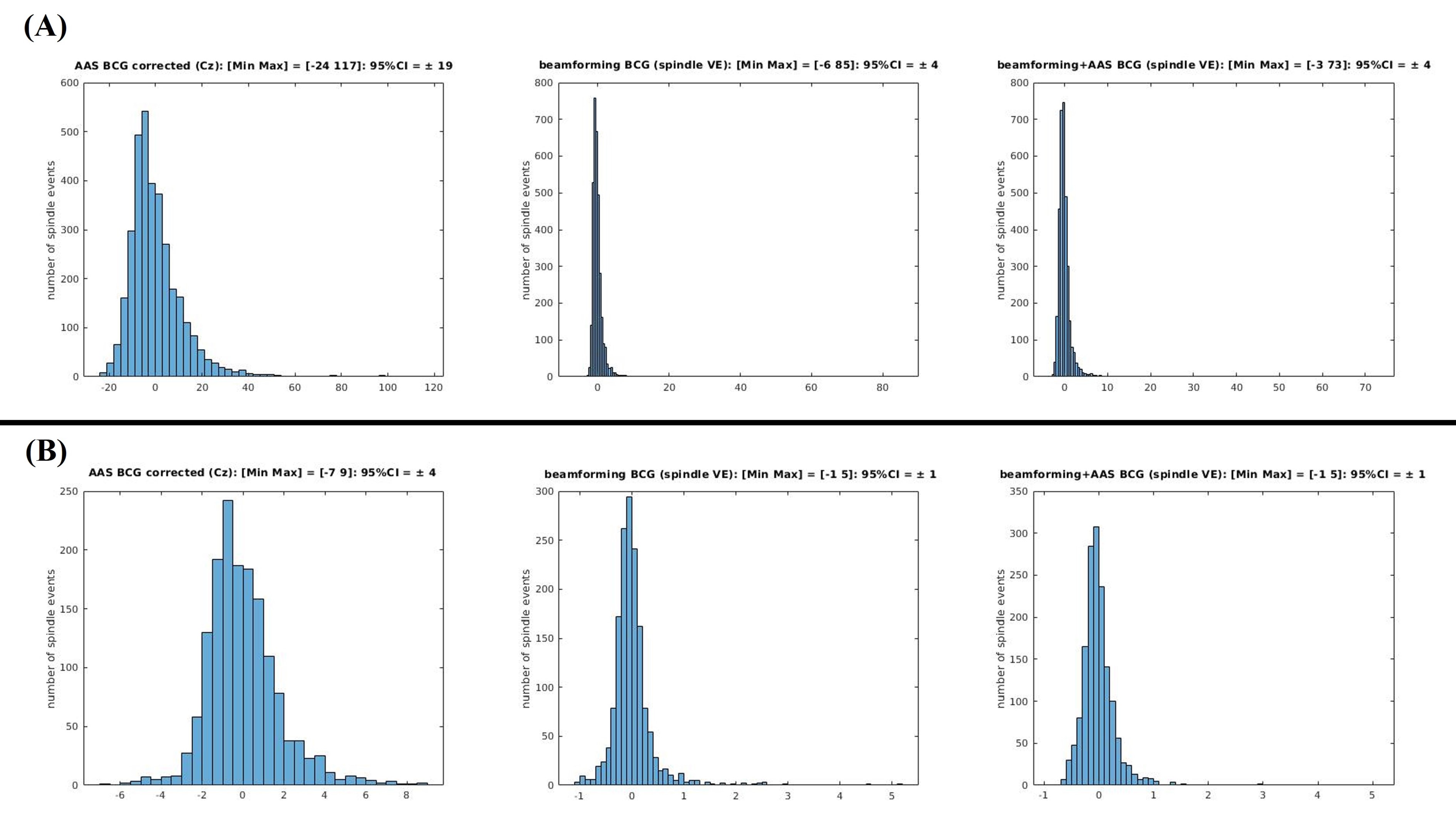

**Figure S9.** Spindle event power amplitude variability across 20 subjects during **(A) the whole nap** and **(B) the validated NREM periods.** Left figures show the spindle amplitude variability from average artefact subtraction (AAS) method ballistocardiogram (BCG) corrected data (Cz: left), middle figures show those from beamforming BCG corrected data (spindle VE: middle), and right figures show beamforming+AAS BCG corrected (spindle VE: right). 95% confident intervals of the channel level spindle power amplitude (AAS BCG corrected) were ±19 (A.U.) during the whole nap and ±4 (A.U.) during the validated NREM periods, whereas those in both source levels (beamforming BCG corrected; beamforming+AAS BCG corrected) were ±4 during the whole nap and ±1 (A.U.) during the validated NREM periods for both source level corrections. These amplitude variabilities were extracted by demeaning each subject’s Hilbert transformed spindle frequency (11-16Hz) signals during each spindle duration. These larger amplitude changes in the channel-level might be due to the residual BCG artefacts causing less accurate spindle power change estimates for the GLM regressors when compared to the source level estimates (GLM3 & GLM4) or just a boxcar ([1 0], GLM1) regressor.
